# Influence of plant host and organ, management strategy, and spore traits on microbiome composition

**DOI:** 10.1101/2020.07.10.178426

**Authors:** Kristi Gdanetz, Zachary Noel, Frances Trail

**Author notes:** corresponding author, 612 Wilson Road, Plant Biology Laboratories, East Lansing, MI, 48824.

## Abstract

Microbiomes from maize and soybean were characterized in a long-term three-crop rotation research site, under four different land management strategies, to begin unraveling the effects of common farming practices on microbial communities. The fungal and bacterial communities of leaves, stems, and roots in host species were characterized across the growing season using amplicon sequencing and compared with the results of a similar study on wheat. Communities differed across hosts, and among plant growth stages and organs, and these effects were most pronounced in the bacterial communities of the wheat and maize phyllosphere. Roots consistently showed the highest number of bacterial OTUs compared to above-ground organs, whereas the alpha diversity of fungi was similar between above- and below-ground organs. Network analyses identified putatively influential members of the microbial communities of the three host plant species. The fungal taxa specific to roots, stems, or leaves were examined to determine if the specificity reflected their life histories based on previous studies. The analysis suggests that fungal spore traits are drivers of organ specificity in the fungal community. Identification of influential taxa in the microbial community and understanding how community structure of specific crop organs is formed, will provide a critical resource for manipulations of microbial communities. The ability to predict how organ specific communities are influenced by spore traits will enhance our ability to introduce them sustainably.

## INTRODUCTION

Maize, soybeans, and wheat are major crops in the United States, with approximately 88, 83, and 47 million acres planted annually, respectively (USDA 2016). Fungal and bacterial diseases are the significant causes of losses in these crop systems (Mueller et al. 2017). In the Midwestern US, the three crops are frequently grown in rotation to preserve soil health and reduce disease. Microbial communities can positively influence multiple plant processes: root-associated microbiota fix nitrogen (Bhuvaneswari et al. 1980; van der Heijden et al. 1998), leaf-associated microbiota play roles in carbon cycling (Abanda-Nkpwatt et al. 2006, Vorholt 2012), and stimulate plant defenses (Vogel et al. 2016). Surveys of microbial communities on agricultural systems have focused on root-associated microbiota (for example Gomes et al. 2018; Rojas et al. 2017; Strom et al. 2019), particularly bacterial communities (for example Mashiane et al. 2017; Rascovan et al. 2016). Studies of fungal communities in row crops have often focused on organ or compartment, most commonly roots and rhizospheres (for example Strom et al. 2019), and less commonly on the leaves and phyllospheres (Karlsson et al. 2014; Karlsson et al. 2017; Sapkota et al. 2017). Several studies have looked at microbial responses to land management or rotation strategies (Karlsson et al. 2017; Longley et al. 2020), but the field sites often are not near replicates, so observed community differences are confounded by divergence in field characteristics (Peiffer et al. 2013; Rascovan et al. 2016). Increased knowledge of the effects of crop rotation on microbiome assembly will inform our ability to manipulate microbiomes for increased yield and disease protection.

Assembly of microbial communities of any plant part results from the distribution of propagules to an appropriate space and their ability to colonize. The resulting fungal community has been shown to be affected by a combination of host genetics (Chen et al. 2020a; Hassani et al. 2018; Maignien et al. 2014) and dispersal, selection and drift of propagules (Vellend 2010, Evans et al. 2016), all influenced by environmental conditions. A large volume of historic literature, focusing mainly on dispersal of ascomycete and oomycete plant pathogens, has shown that spore morphology (size, shape, melanization) and dispersal mechanisms (active dispersal, passive dispersal including hypogeous dispersal, and motility) play a critical role in the distribution of fungal propagules (Aylor 2017; Calhim et al., 2018; reviewed by Deacon 2006; Schumann & D’Arcy 2010). Other studies have focused on the dispersal of basidiospores along with the interactions of spore traits with the environment (Calhim et al., 2018; Halbwachs et al. 2015; Peay et al. 2012; Pringle et al. 2015).

Compared to fungi, bacterial dispersal mechanisms are less varied due to less diverse morphology and modes of distribution. Plant-associated bacteria are presumed to predominantly colonize from the soil (Compant et al. 2010). Chemotaxis driven by bacterial flagella and quorum sensing can affect colonization of the plant surface above- and below-ground. Migration to above-ground parts can also occur through endophytic colonization of the vascular system (Lamb et al. 1996). Bacteria are distributed in hotspots across the surface of the above-ground plant parts (Lindow & Brandl 2003) with the bulk of the colonization on lower leaf surfaces (Remus-Emsermann et al. 2014). Furthermore, bacteria can disperse along fungal hyphae, demonstrating the importance of bacterial-fungal interactions for the assembly of the microbiome (Kohlmeir et al. 2005; Zhang et al. 2018).

In the present study, we aimed to conduct a comprehensive analysis of the fungal and bacterial communities of wheat, maize, and soybeans across a full three-year rotation cycle under different land management strategies. Previous studies at the same site characterized wheat and soybean microbiomes within single growing seasons (Gdanetz & Trail 2017; Longley et al. 2020). We investigated the microbial communities associated with each of the plant organs (leaf, stem, root) and the change in communities across growth stages. To achieve this goal, we analyzed the fungal and bacterial communities of one-hectare replicate plots in a completely randomized block design managed as organic, conventional till, no-till, and low input for over thirty years (Robertson 2015). We compared these microbial communities to our previous studies of the wheat microbial community at the same site. Our analysis of organ associated microbiomes revealed fungal taxa that are specific to a single plant organ and demonstrates the role of dispersal mechanisms and spore traits associated with these different niches resulting in the specificity of fungi to above- or below-ground plant organs.

## MATERIALS AND METHODS

### Site Description and Sample Collection

The Michigan State University W. K. Kellogg Biological Station Long-Term Ecological Research site (KBS-LTER) is located in Hickory Corners, Michigan, USA (42.411085, −85.377078). The KBS-LTER Main Cropping System Experiment site has been planted in a wheat-maize-soybean rotation series since 1993, and is organized in randomized, replicated plots under four land management strategies with six replicate plots for each strategy (24 total plots): T1 = conventional till, T2 = no-till, T3 = reduced chemical inputs with red clover cover crop, T4 = organic with red clover cover crop (Robertson 2015). Average annual precipitation for this site is 1,005 mm, with approximately half falling as snow (NCDC 1980-2010 climate normals for the Gull Lake Biological Station, https://lter.kbs.msu.edu/research/site-description-and-maps/general-description/climate-normals). The annual precipitation was 1177 mm, 932 mm, and 1153 mm during wheat, maize, and soybean growing years, respectively. Detailed site and management information can be obtained from https://lter.kbs.msu.edu/research/.

For the 2013 harvest, wheat seeds of the cultivar, Pioneer 25R39 soft red winter wheat (Pioneer High-Bred International, Inc., Johnston, IA, USA), were planted in all plots, but wheat seeds for conventional, no-till, and low inputs plots were coated with Gaucho insecticide at purchase (Bayer Crop Science, Research Triangle Park, NC, USA). These have been previously analyzed and reported (Gdanetz & Trail 2017), but the data were used here as a comparison for the three-crop rotation. For the 2014 harvest, a commercial maize hybrid cultivar, Dekalb DKC52-59 maize hybrid (Monsanto Company, St. Louis, MO, USA), was planted in plots T1-T3; and an organic-approved cultivar, Blue River Hybrids 25M75 organic maize (Blue River Organic Seed, Ames, IA, USA), was planted in T4 plots. For the 2015 harvest, the commercial soybean cultivar, Pioneer P22T69R (Roundup Ready®) soybean seed (Pioneer High-Bred International, Inc., Johnston, IA, USA), was planted in T1-T3 plots; and Viking Organic Soybean Seed, Variety 0.2265 (Albert Lea Seed, Albert Lea, MN, USA), was planted in T4 plots. The conventional, no-till, and low input management used fungicide- and insecticide-treated maize and soybean seed. Plants were collected at three analogous developmental stages for each host species: late vegetative growth, inflorescence, and early seed/ear/pod development (Table S1). At each developmental stage, three intact plants were removed from each of the 24 plots. Fine and thick roots, and above-ground organs (leaves and stems) from each plant were placed in separate sterile sample collection bags (Nasco Whirl-Pak®, Fort Atkinson, WI, USA) and maintained on ice during transport. Roots were rinsed to remove loosely attached soil, and plants were stored at −80°C, then lyophilized. Lyophilized plants were stored at room temperature under a desiccant until grinding and DNA extraction.

### Sample and Sequence Processing

Lyophilized plants were separated by organ (leaf, stem, root) before grinding with a Retsch Oscillating Mill M400 (Verder Scientific, Newtown, PA, USA) and DNA was extracted using the Mag-Bind® Plant DNA Plus Kit (Omega Bio-tek, Norcross, GA, USA) following the manufacturer’s protocols with the KingFisher™ Flex (ThermoFisher Scientific, Waltham, MA, USA). Fungal ITS2 and bacterial 16S rRNA gene libraries were generated as described previously and sequenced using Illumina MiSeq 2×250 bp chemistry (Table S1) (Gdanetz & Trail 2017; Kozich et al. 2013; Toju et al. 2012). Sequences are available with the National Center for Biotechnology Information Small Read Archive project. Accession numbers for wheat, maize, and soybeans are SRP102192, SRP102245, and SRP120500, respectively.

Forward and reverse read pairs from each library were merged with USEARCH (version v8.1.1861) (Edgar 2010; Edgar et al. 2011; Edgar & Flyvbjerg 2015). Low quality reads, (fastq expected error set to 1.0), and read pairs without mates were discarded (read length was set at 250 bp for 16S and 380 bp for ITS2 sequences). The 16S and ITS2 libraries from all three host species were concatenated into one large dataset for each barcode region. Sequences were processed as described previously (Gdanetz & Trail 2017). Briefly, the USEARCH pipeline was implemented for chimera filtering and OTU assignment, with the cluster threshold set to 97% similarity. The Ribosomal Database Project Classifier was used for taxonomic assignment, with the 16S and the UNITE ITS training sets for bacteria and fungi, respectively, customized to include the plant hosts (Cole et al. 2013; Deshpande et al. 2016; Wang et al. 2007). OTUs matching plants, mitochondria, and chloroplasts, unidentified at the Kingdom or Domain level, or matching non-target organisms were discarded. Samples were filtered to include OTUs that occurred in at least five samples and this trimmed dataset was used for all downstream analyses.

### Analysis of Organ-Specific OTUs

Organ-specific fungal OTUs were identified during core OTU analysis. To reduce the number of OTUs that were associated with an organ due to chance, such as a bloom of fruiting bodies near plant collection sites, organ-specific lists were further filtered to include OTUs found in a minimum of five replicate plots at the field site. Initial examination of the fungal OTUs using FUNGuild (Nguyen et al. 2016) suggested that there was specificity of taxa to plant organs. To determine if the organ colonization of fungal OTUs was due to differences in dispersal mechanisms across lineages of fungi, propagule characteristics were defined for organ-specific OTUs with taxonomic assignments through primary literature when available (Table S6 and references therein).

Propagule characteristics were classified into propagule size (length on longest side), cell motility, presence of yeast forms, melanization, and means of dispersal. For spore size, we chose three categories (up to 10 µm, 10-100 µm, and greater than 100 µm) based upon the natural breaks in spore size occurring between higher taxonomic ranks. Motile spores and basidiospores were not placed in size groups, but formed separate groups due to their special modes of dispersal in the former, and the unique size constraints in the latter which are dictated by size limitations and fruiting body structure (Galente et al. 2011; Fischer et al. 2010; Stolze-Rybczynsk et al. 2009). Pigmentation is also characteristic of some types of spores, thus melanization status was included when known. Dispersal mechanisms included aerial dispersal, water dispersal (due to rain, pools, or flowing water), soil dispersal and hypogeous. Finally, yeasts are often present on plant surfaces, and we distinguished those species that are often in a yeast form. The category “other” was composed of fungi which had less than 5 representative significant OTUs in the indicator species analysis or categories with less than than 0.2% mean relative abundance, which included: spores up to 10 µm dispersed in the soil or aerially by water; spores up to 10 µm melanized and dispersed aerially; spores up to 10 µm hypogeous; spores 10 to 100 µm dispersed by insects, soil, water, or were hypogeous; spores greater than 100 µm dispersed by water or aerially; and yeasts dispersed aerially or by water. Propagule characteristics assigned to OTUs can be found in Table S6 and references therein.

After classification of fungal propagule characteristics, an indicator species analysis was conducted to determine which fungal OTUs were significantly associated with a host organ. Samples with less than 1000 reads were removed, then read counts were normalized using the cumulative sum scaling technique with ‘metagenomeSeq’ package (Paulson et al. 2013). Indicator species analysis was performed with the R package ‘indicspecies’ (version 1.7.6) to identify OTUs significantly associated with host organs (De Caceres & Legendre 2009). Rare taxa with a mean relative abundance below 10^−5^ within each host species were not included in the analysis and a Benjamini-Hochberg p-value correction was used for multiple testing. Following indicator species analysis, the compositional abundance of each OTU was plotted in a ternary diagram using ‘ggtern’ (Hamilton & Ferry 2018) and colored by propagule characteristic categories.

### Community Analysis

Venn diagrams and tables for core taxa analysis were generated using the gplots package (Warnes et al. 2016) in the R statistical computing environment (version 3.3.3). Before generating bar plots, taxa were merged at class level. Taxa that were present in less than 3% of the samples were removed. Relative abundances of taxa were calculated as a percentage of total sequences in each sample. OTUs per sample were normalized with variance-stabilizing normalization and Bonferroni-corrected p-values were used to identify significantly different OTUs across samples with the ‘DESeq2’ package (McMurdie & Holmes 2014).

To determine the effect of management strategy on microbial community diversity, comparison between host species, organs within and between each growth stage, and comparisons between management were made for each host. Alpha diversity statistics (abundance transformation, observed OTUs, and Shannon’s Index) were calculated with the ‘phyloseq’ package (McMurdie & Holmes 2013). Non-rarefied data were used to calculate the Shannon Diversity Index (H’), an alpha diversity metric that measures species diversity within a sample (McMurdie & Holmes 2014), and analysis of variance (ANOVA) and Tukey’s honest significant difference test (HSD) were used to determine significance. Non-metric multidimensional scaling (NMDS) ordination analysis was conducted using Bray-Curtis distance values, a measure of species diversity between communities. Permutational analysis of variance (PERMANOVA) tests were completed using the adonis function in the ‘vegan’ R package (Oksanen et al. 2016). Heterogeneity of the variances was calculated using the betadispersion function of the ‘vegan’ package (Oksanen et al. 2016). Figures were generated with the ‘ggplot2’ package (Wickham 2009).

Putative hub taxa, species that influence the presence of others in an environment, were identified as highly connected nodes based on calculations by Agler et al. Fungal and bacterial OTU tables were concatenated before calculating the co-occurrence network, leaf- and stem-specific OTUs were grouped together as phyllosphere samples and filtered to include OTUs with ≥20 reads in ≥10 samples using the SpiecEasi package in R (Kurtz et al. 2015). Network statistics and network plots were generated using Cytoscape (version 3.5.1, http://www.cytoscape.org). Network statistics, specifically log-transformed degree (number of connections between nodes), betweenness centrality (measure of nodes with shortest connections to other nodes), and closeness centrality (measures distances between nodes), were fit to a normal distribution, and OTUs in the 75th and 95th percentile across all three statistics were identified as putative hub taxa.

## RESULTS

### Observed and Core Taxa

To investigate the effect of crop management strategy (conventional, no-till, low input, organic) on microbiomes of leaves, stems, and roots, samples were collected across replicate plots throughout the season at three growth stages (Table S2). We analyzed bacterial and fungal communities of maize and soybeans, grown under the four management strategies over a two-year period (maize in 2014 and soybeans in 2015), using fungal (ITS2) and bacterial (16S) rRNA gene amplicon sequencing. By including sequencing for wheat from the 2013 field season (Gdanetz & Trail 2017), we analyzed one full rotation cycle from this field site. Following filtering and quality control, high quality reads were identified for fungi and bacteria, respectively, with means of 21,245 fungal and 42,166 bacterial reads per sample (Table S3). No fungal sequences from soybean T4 stems and roots passed quality control (Table S4) and thus were not included in the rest of the analysis. A total of 4,739 fungal and 8,942 bacterial high quality OTUs were identified for maize and soybean combined. Excluding off-target kingdom and domain matches, the combined data from all three crops revealed 7,233 fungal and 12,673 bacterial OTUs (Table S3).

Unique and shared OTUs were compared among the three host species and plant organs (leaves, stems, and roots). Of the bacterial and fungal OTUs, fewer OTUs were unique to soybean plants (Fig. 1, 2). Bacterial communities were similar across all management strategies within a host species at the Class level (Fig. S1), however we observed differences among hosts and organs (Fig. 1). Fungal OTUs were more uniformly distributed across all hosts and management strategies than bacteria.

**Figure 1.**
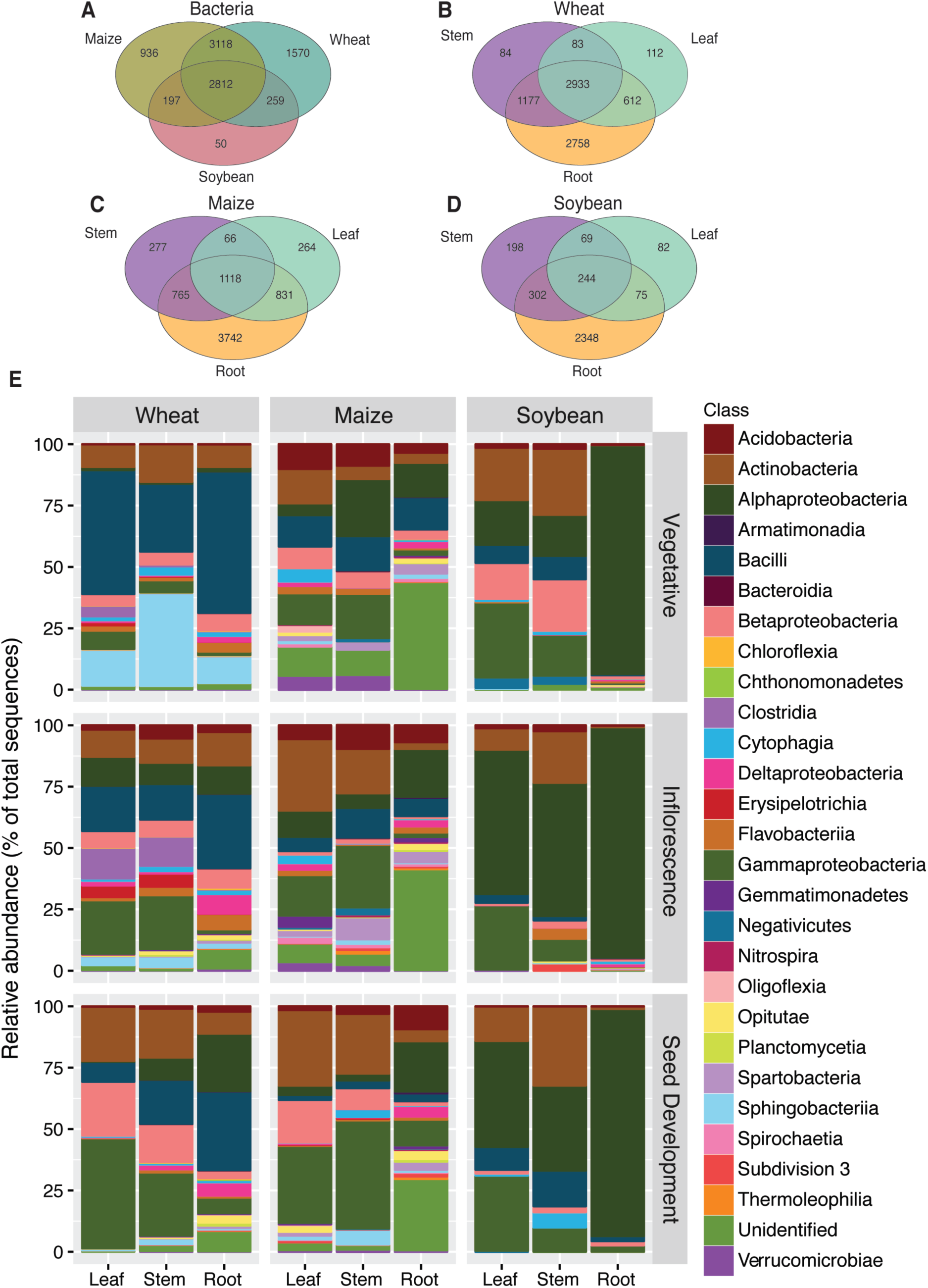
Distribution of OTUs across crops and organs. Shared and unique bacterial operational taxonomic units across (A) whole plants and (B, C, D) organs in wheat, maize, and soybean respectively. (E) Class-level relative abundance of operational taxonomic units in bacterial communities across crop, organ, and growth stage.

**Figure 2.**
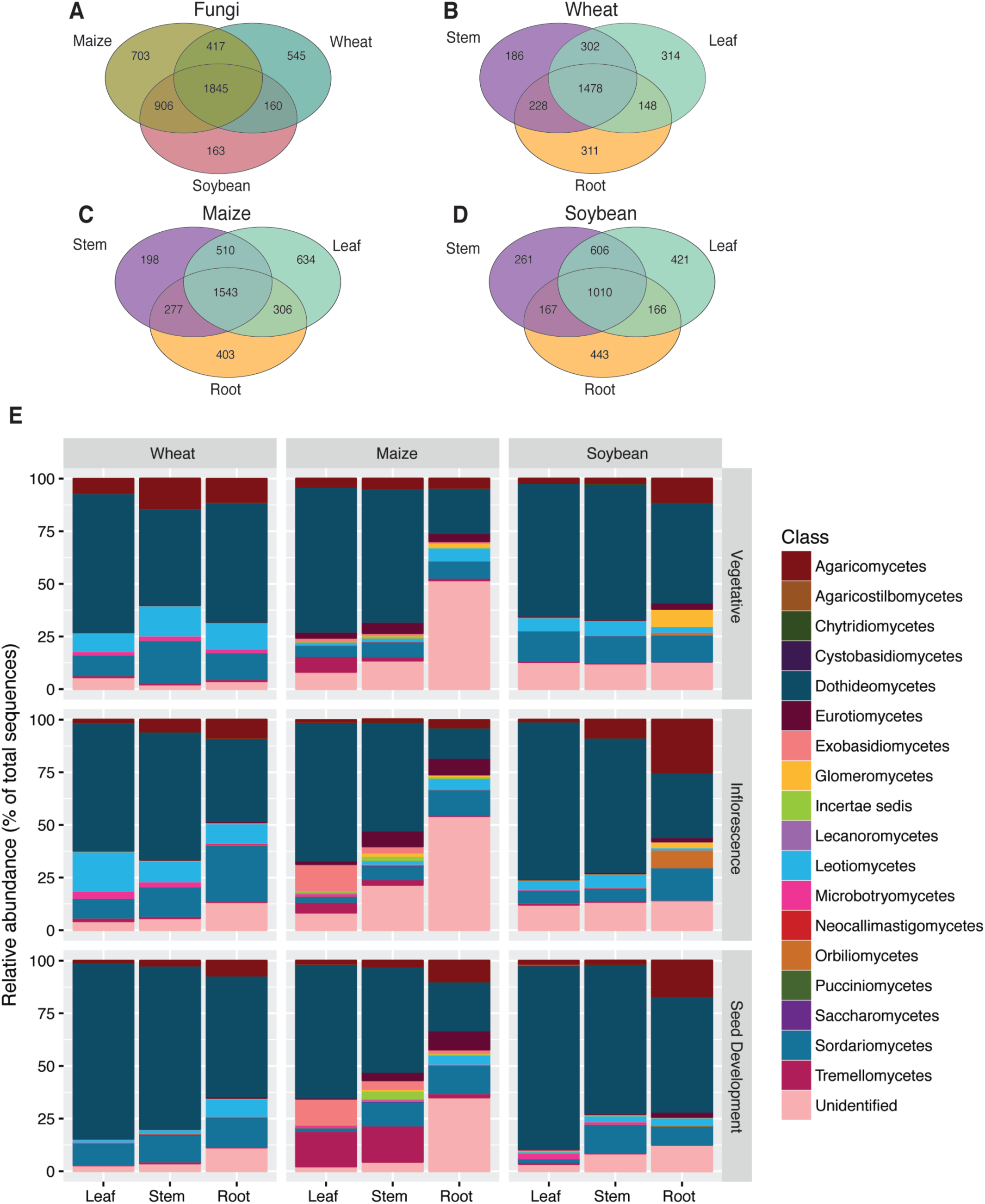
Distribution of OTUs across crops and organs. Shared and unique fungal operational taxonomic units across (A) whole plants and (B, C, D) organs in wheat, maize, and soybean respectively. (E) Class-level relative abundance of operational taxonomic units in fungal communities across crop, organ, and growth stage.

Highly-abundant bacterial Classes Actinobacteria, Alphaproteobacteria, Bacilli, Betaproteobacteria, and Gammaproteobacteria were generally shared across all host species (Figs. 1E, S1). Maize and soybean bacterial OTUs showed organ-specificity, similar to those observed for the 2013 wheat analysis (Figs. 1B, 1E; Gdanetz & Trail 2017). Roots consistently had the most organ-specific bacterial OTUs, with higher numbers of bacterial OTUs unique to roots than shared among organs in either maize or soybeans (Fig. 1). Soybean roots contained 88% of the host-specific bacterial taxa, compared with 93% in wheat and 87% in maize (Fig. 1). In maize roots, 246 OTUs were significantly different in abundance across management strategies and growth stages (p<0.01, Table S5). In total, 255 bacterial OTUs from roots were identified as significantly different across management strategy and growth stage (p<0.01, Table S5). The phyllosphere organs, leaves and stems, of each host revealed similar Classes of bacterial taxa when compared with the roots of the same host species (Figs. 1, S1). Bacterial community composition of wheat and maize above-ground organs changed with growth stage (Fig. 1E). OTUs found in maize stems were more consistent between no-till and low input samples than other management strategies (Fig S1). The dominance of Alphaproteobacteria in soybean roots of all management strategies was not surprising since this Class includes Rhizobia.

At the OTU level, 266 fungal OTUs (5% total fungal OTUs) were identified as significantly different (p<0.05) across management strategies and growth stages (Table S5). The Classes Agaricomycetes, Dothideomycetes, Leotiomycetes, and Sordariomycetes occupied the highest percentages of OTUs for each of the crop organs and growth stages (Fig. 2). Most notable were the Dothidiomycetes, the largest class in the kingdom Fungi and common plant pathogens and melanized endophytes, which dominated all growth stages and treatments (Fig. 2, S2). Also notable is the presence of Microbotryomycetes predominantly on wheat in the two earlier stages. These members of the Pucciniomycotina often present as epiphytic yeasts. Yeasts that inhabit the phyllosphere are well suited to the oligotrophic conditions present on leaf surfaces. They produce extracellular polysaccharides, surfactants, and carotenoid compounds, which may be important for maintaining biofilms and stress tolerance (Cobban *et al*., 2016; Fonseca and Inacio 2006)..

Fungal communities in roots of maize and soybeans differed from those found on above-ground organs (Fig. 2). Root communities in wheat also differed from above-ground organs, but these observations were dependent on plant age (Fig. 2); above-ground communities of wheat were more similar during vegetative growth (Fig. 2; Gdanetz & Trail 2017). Root samples of all hosts and growth stages, except the wheat vegetative stage, had a higher abundance of Agaricomycetes than did the phyllosphere organs (Figs. 2, S2). The Basidiomycota Class Tremellomycetes, which includes plant-associated yeast forms, were highly represented in phyllosphere organs on maize, but at very low levels in other samples (Figs. 2, S2). The Glomeromycetes (the group containing the arbuscular mycorrhizal fungi) and Orbiliomycetes (nematode trapping fungi) were more abundant in soybean roots than other host plant species (Figs. 2, S2).

### Analysis of abundance of single OTUs

The most abundant fungal order present was the Pleosporales in the phyllosphere with 413 OTUs present across all crops. The Glomerales in the roots are the next most abundant order with 109 OTUs. OTUs of the Agaricales, Capnodiales, Chaetothryiales, Helotiales, Hypocreales, and Sordariales are the next most abundant orders, with each of these containing 30 to 70 unique OTUs. The Sordariales (Sordariomycetes) are predominantly associated with roots.

The most abundant OTU was OTU_2, *Epicocum nigrum*, and was specific to leaves: maize (R.A. 13.7%), soybean (R.A. 5.82%), wheat (R.A. 1.57%). OTU_413 and OTU_804 were found abundantly on soybean stems (R.A. 8.29%, 3.08%), maize stems (OTU_804 R.A. 1.55%), and on leaves of wheat (OTU_413 R.A. 3.60%; Table S6). These OTUs were assigned to *Stagonosporopsis loticola* (Chen & Kirschner 2018), a lotus pathogen, but are likely to be related members of the Didymellaceae. OTU_8 (an unidentified basidiomycete) was the second most abundant OTU and was found on maize leaves (R.A. 11.16%) and on soybean roots (R.A. 0.53%). Individual OTUs with high relative abundance on roots were mainly assigned to order or family, which obscured the specificity of their life history. Interestingly, all seven of the OTUs significantly associated with the stem fell into the Chytridiomycota. We have assigned these OTUs to the category “zoospores/water” in keeping with the major phenotypes of that phylum (James et al. 2006). None of these OTUs were defined at lower taxonomic ranks and although three were associated with wheat, three with soybean and one with maize, they represented seven separately defined OTUs.

### Organ-Specific OTUs

Analyses of OTUs conducted using higher taxonomic classifications such as in Fig. 1 do not supply us with adequate information on the biology of the fungi, therefore we examined the identity of unique fungal OTUs for each plant organ across all host species (Fig. 3a-b; Table 1), and determined their spore traits and dispersal mechanisms from the literature. Our objective was to evaluate the relationship between fungal distribution and organ specificity. Although many OTUs were shared across organs (Figs. 2a-d, 3a-b), there was identifiable organ-specificity among the plant-associated fungal taxa. Across all hosts, 2591 root-, 112 stem-, and 2257 leaf-specific fungal OTUs were identified. A total of 1506 root-, 112 stem-, and 1789-leaf specific OTUs were assigned a taxonomic classification specific enough to determine their spore traits and dispersal mechanisms (Table S6); the remaining OTUs could not be assigned spore categories. Categories of propagule characteristics were defined as spore size and type, motility and predicted means of dispersal based on the literature. Distribution of propagule characteristics were analyzed to determine if location of organ specific taxa aligned with their dispersal and other spore characteristics. Spores from 10 to 100 µm comprised 57% of the spores of all taxa identified, and 71% of leaf-specific taxa, 64% of stem-specific taxa, and 41% of root-specific taxa; the majority of taxa from each organ produced predominantly melanized spores and were at least in part aerially distributed. These melanized, aerially distributed taxa together with the taxa producing spherical spores (mainly very large) associated with the Glomeromycota (28% of taxa identified as associated with roots), formed the predominant root-specific fungi. Taxa forming spores “up to 10 µm” represented 9% of the leaf-specific taxa, 11% of the stem-specific taxa, and 13% of the root-specific taxa and fell into several different categories of dispersal, with the largest numbers of OTUs in “aerial, soil”, “hypogeous”, “insect, other”, and the “up to 10 um, lichenized/aerial, soil”. Those spores in “aerial, soil” dispersal categories demonstrated a higher distribution to above ground versus below-ground (74% versus 26% in “aerial, soil”, and 58% versus 42% in the “lichenized/aerial, soil”). Those OTUs in the “hypogeous” and “insect, other” categories showed exclusive distribution to the soil (Table S6). Other dispersal categories within the “up to 10 µm” group had very low membership.

**Table 1.** Counts of the number of fungal OTUs significantly associated with leaves, stems, or roots across wheat, corn, and soybeans.

| **Spore Category** | **Leaf** | **Stem** | **Root** | **Total** | **Mean % RA** |
| --- | --- | --- | --- | --- | --- |
| 10 to 100 microns, usu. melanized / aerial, water | 206 | 35 | 102 | 343 | 14.2 |
| basidiospores / aerial | 31 | 6 | 24 | 61 | 2.1 |
| yeast / other, soil | 20 | 4 | 4 | 28 | 2.0 |
| other<sup>a</sup> | 5 | 10 | 17 | 32 | 1.5 |
| 10 to 100 microns / aerial, water | 10 | 10 | 9 | 29 | 1.4 |
| up to 10 microns / aerial, soil | 13 | 2 | 5 | 20 | 0.9 |
| AM fungi 50 microns to 1 mm / hypogeous | 2 | 6 | 81 | 89 | 0.6 |
| other / soil | 0 | 0 | 3 | 3 | 0.5 |
| up to 10 microns, melanized / soil | 0 | 1 | 1 | 2 | 0.3 |
| zoospores / water | 0 | 1 | 15 | 16 | 0.04 |
| up to 10 microns / hypogeous, aerial | 0 | 0 | 13 | 13 | 0.03 |
| up to 10 microns, lichenized / aerial, soil | 7 | 0 | 5 | 12 | 0.03 |
| up to 10 microns / insect, other | 0 | 0 | 10 | 10 | 0.03 |
| basidiospores, yeasts / aerial, other, water | 2 | 1 | 3 | 6 | 0.02 |

<sup>a</sup>Other category includes spore categories with fewer than five representative significant OTUs, or categories with mean percent relative abundance was less than 0.2

**Figure 3.**
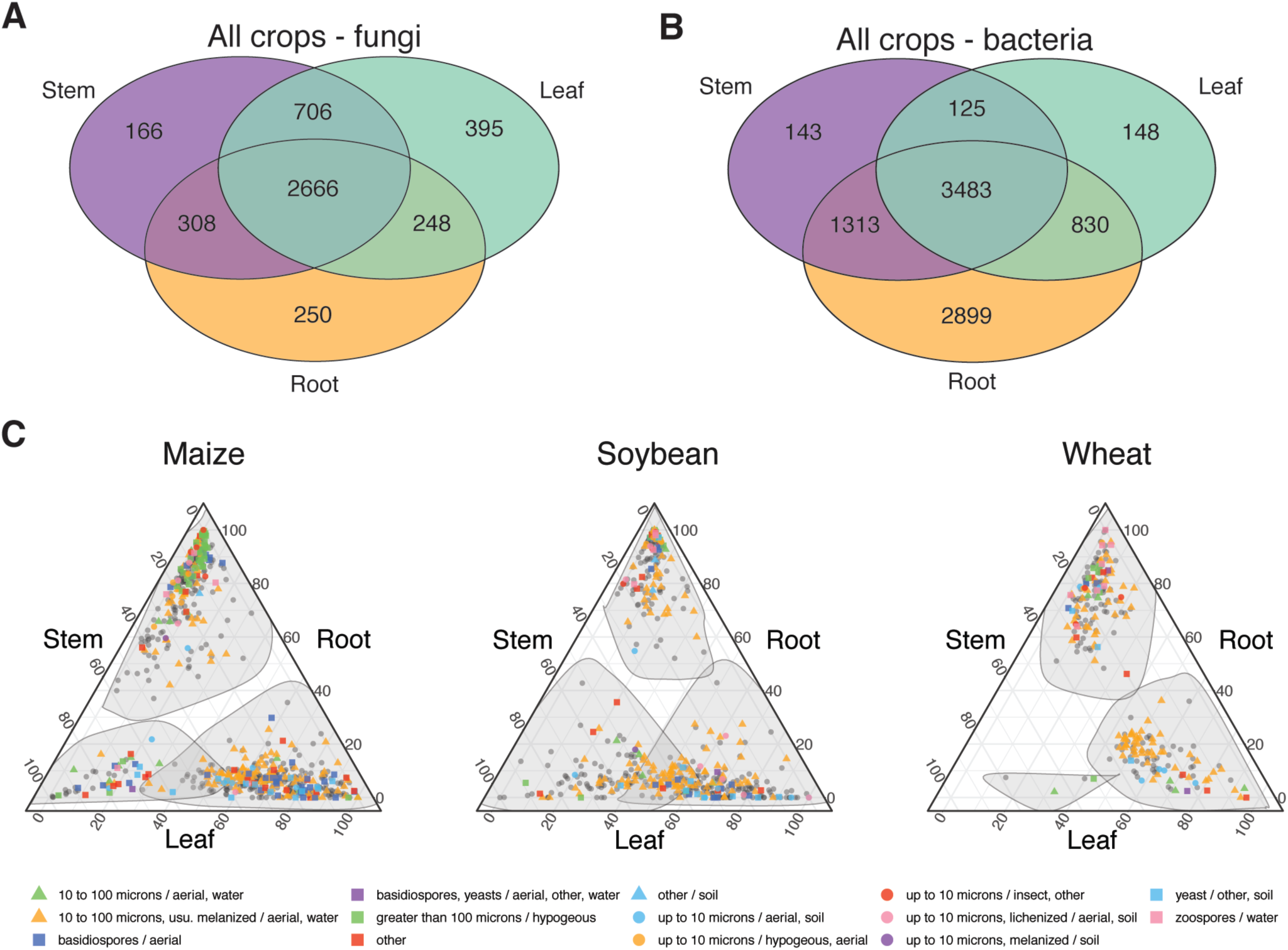
Microbes associated with plant organs. (A) Fungal and (B) bacterial shared and unique OTUs among plant organs. Additional information on spore dispersal and function can be found in Table S5. (C) Ternary plots illustrating the composition of each fungal OTU in each plant organ-crop combination. Colored points were significant indicators of a particular plant organ with a Benjamini-Hochberg corrected P-value (α = 0.05). Shaded areas encircle the OTUs that are significant indicators for a plant organ. Points are colored by spore category to illustrate the distribution of spore categories within each plant organ. The distribution of spore categories across crops is shown in Fig. S4.

To visualize the data from the three organs, we used indicator species analysis combined with ternary plots for illustration of the distribution of spore categories for each plant host (Fig. 3c, Table S6) and for representation of the distribution of each of the categories among all hosts (Fig. S3). The results demonstrated that the microbiomes of maize, soybean, and wheat were distinct between organs. Taxa forming spores up to 10 µm in diameter were slightly more likely to be in leaves if aerially distributed and found in soil (13 leaf, 2 stem, 5 root OTUs), but if aerial and hypogeous, or dispersed by insects, were exclusively associated with roots (13 hypogeous root OTUs; 10 insect dispersed root OTUs; Table 1). The taxa in the Glomeromycota, which produces very large hypogeous spores, were mostly associated with maize roots (33 OTUs) and soybeans (43 OTUs), and were highly reduced in wheat roots (5 OTUs) (Table S6). Yeasts that were in the Basidiomycota were more likely to be associated with leaves (20 leaf, 4 stem, 4 root OTUs; yeast / other, soil; Table 1). Both roots and leaves were populated with aerially distributed, melanized taxa in the size range of 10-100 µm (206 leaf, 35 stem, 102 root OTUs; Table 1). The zoospore forming taxa were almost exclusively in roots (0 leaf, 1 stem, 15 root OTUs), with wheat harbored the majority (50%) out of all the host plants (Table S6). Maize hosted the greatest numbers of leaf-specific basidiospore producing taxa. Finally, maize had the most discrete stem-specific taxa and was colonized by more yeast-producing taxa on the leaves and stems than the other plant hosts.

### Community Diversity

Conventional and transgenic cultivars, in maize and soybeans, did not differ significantly in alpha diversity (as measured by *H*’) of fungal and bacterial communities when comparing the same growth stage and plant organ (Fig. S4, Tables S7-S8). The effect of cover crop presence or absence, in low input and organic plots, on fungal and bacterial communities inhabiting each of the organs (leaves, stems, or roots) was compared for each collection period and host species. *H’* of fungal communities were significantly different between plots with and without cover crops in soybean stems during the seed development stage only (Fig. S5, Tables S7-S8). *H’* of root bacterial communities during flowering of wheat was significantly lower in plots with cover crops (p < 0.001; Fig. S5, Tables S7-S8). There were no other observed differences in alpha diversity between hosts with and without cover crops (Fig. S5, Tables S7-S8).

The effect of management strategies on the microbial community within organs was assessed. Overall, bacterial alpha diversity of wheat, maize, and soybean roots was significantly higher than *H’* of leaves and stems of these host species, but there were not significant differences in *H’* of fungal communities between crop organs (Figs. 4c-d, Tables S7-S8). Bacterial *H’* of maize roots was higher than that of above-ground organs at all growth stages and under all management strategies, this was a significant increase except in leaves and stems of conventional, no-till, and low input of maize at the vegetative stage (Figs. 4c, S6b, Tables S7-S8). The bacterial diversity of soybean roots was more variable than maize and decreased across the growing season (Figs. 4d, S6b). Bacterial communities of soybean leaves and stems during pod development from all management strategies had significantly higher *H’* compared to roots, except stems from the organic management strategy, due to the dominant presence of Rhizobia OTUs (Tables S7-S8). The mean fungal diversity of maize roots trended lower than maize above-ground organs at the vegetative stage, but was significantly lower in all managements when compared with fungal diversity of maize leaves from conventional plots (Fig. 4a, Tables S7-S8). Whereas fungal diversity in soybean roots decreased as plants aged, this trend was only significant within the low input plots (Fig. 4b, Tables S7-S8).

**Figure 4.**
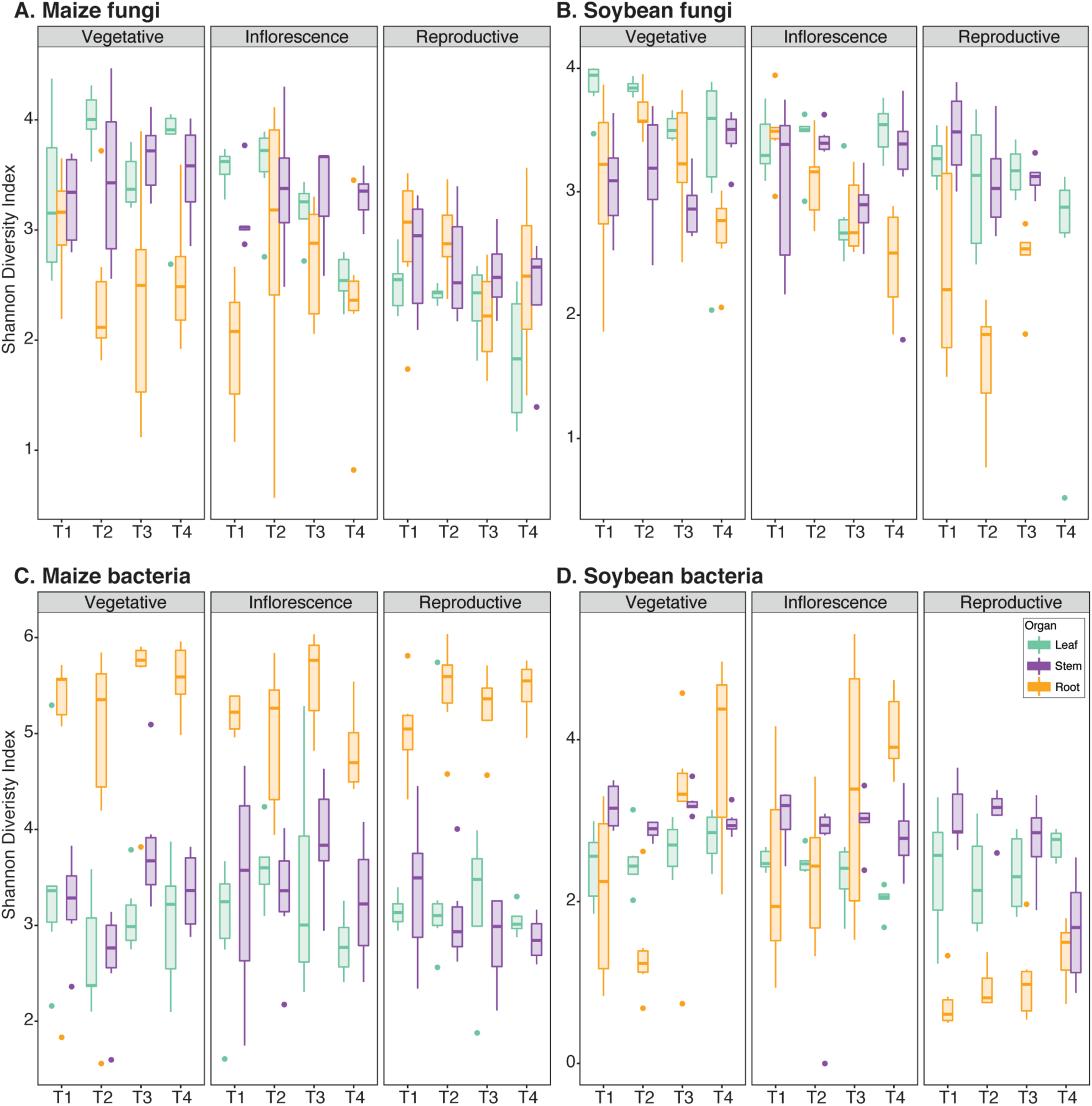
Alpha diversity maize fungi (A), soybean fungi (B), maize bacteria (C), soybean bacteria (D) found on different plant organs across growth stages, and under four different management strategies, estimated by Shannon Diversity Index. T1=conventional, T2=no till, T3=low input, T4=organic. Reproductive growth stage indicates early ear development, and early pod fill for maize and soybean, respectively. Data are represented by six replicates from each stage-management-organ combination. Center line of boxes represents median of samples. The upper and lower sides of the boxes represent the third and first quartiles, respectively. Whiskers represent ± 1.5 times the interquartile range. Data points beyond whiskers represent outliers. Analysis of variance and Tukey’s honest significant difference were used to test significance (P < 0.05). Statistical support is detailed in Table S5.

Beta diversity of microbial communities was visualized with NMDS plots generated from Bray-Curtis distances (Figs. 5, S7). Global analysis revealed clusters for each host species and segregation was clear for above-ground versus below-ground samples in fungi (Figs. 5a, S7a, S7c). At the vegetative growth stage, wheat bacterial communities from all samples formed a clearly defined cluster, while maize and soybean communities shared a segregation between above- and below-ground; however, the separation of wheat from the other host species was lost in the later growth stages (Fig. 5b, S8). Within maize (Figs. S7a-b) and soybean communities (Figs. S7c-d), there was clustering of fungal and bacterial communities by plant organs. Centroids of clusters were significantly distinct as determined by PERMANOVA, and variances were also significantly different (*P* < 0.001, Table S9). Within each growth stage, there was not clear clustering of fungal and bacterial communities by land management strategy; although, root samples formed clusters separate from above-ground organs (Fig. S8). NMDS plots of management strategies had significant centroids (p < 0.01); significantly different variances were observed for fungal communities (p < 0.05), but were not significant for bacterial communities (Table S9). Ordination analysis of the effect of cover crop or cultivar on microbial communities, showed similar patterns to ordination analysis of management strategy; points segregated by above- and below-ground samples (Figs. S9-S10, Table S9).

**Figure 5.**
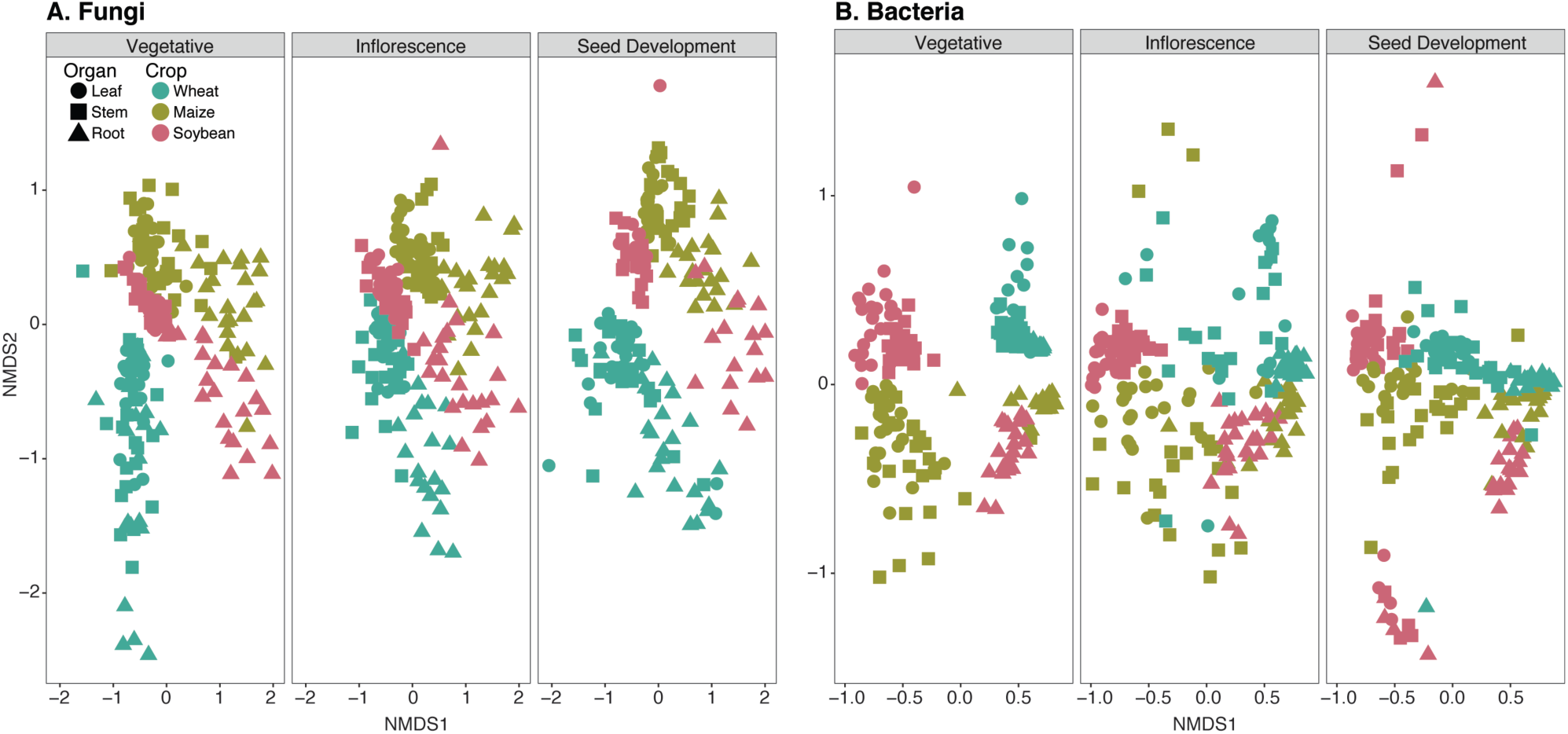
Effect of growth stage on beta diversity of fungal (A) and bacterial (B) communities on all crops. Points colored by crop species, shapes indicate plant organ. Non-metric multidimensional scaling (NMDS) calculated by Bray-Curtis distance. Statistical support is detailed in Table S9.

### Microbial Networks

Networks were generated with OTUs from all management strategies, and growth stages. Networks from individual growth stages of one host did not meet filtering requirements, therefore all growth stages were analyzed together. Microbial hubs within networks were defined as nodes with a high degree of connectivity, betweenness centrality, and closeness centrality (Agler et al. 2016) and were composed of species that were likely to influence the presence of others in the environment. Across all three host species, 111 putative hubs were identified and, using more stringent criteria (95^th^ versus 75^th^ percentiles), the number of hub taxa was reduced to 17 (Table 2). Of the putative hub taxa, nine bacterial hubs were shared between wheat roots and the phyllosphere (Table 2, Figs. 6, S11). All hub taxa in wheat roots were correlated with at least one other hub taxon in the network, and six of these hubs had at least one negative correlation with another node (Fig. 6). *Metarhizium* was the only hub taxon in the wheat phyllosphere not correlated with other hub taxa (Fig. 6). Of the wheat phyllosphere hub species, *Acidovorax* sp., *Discula destructiva, Helvella dovrensis, Luteolibacter* sp., *Metarhizium* sp., *Methylophilus* sp., *Pedobacter* sp., and *Taibaiella* sp. did not have any negative co-occurrence (Fig. 6).

**Table 2.**
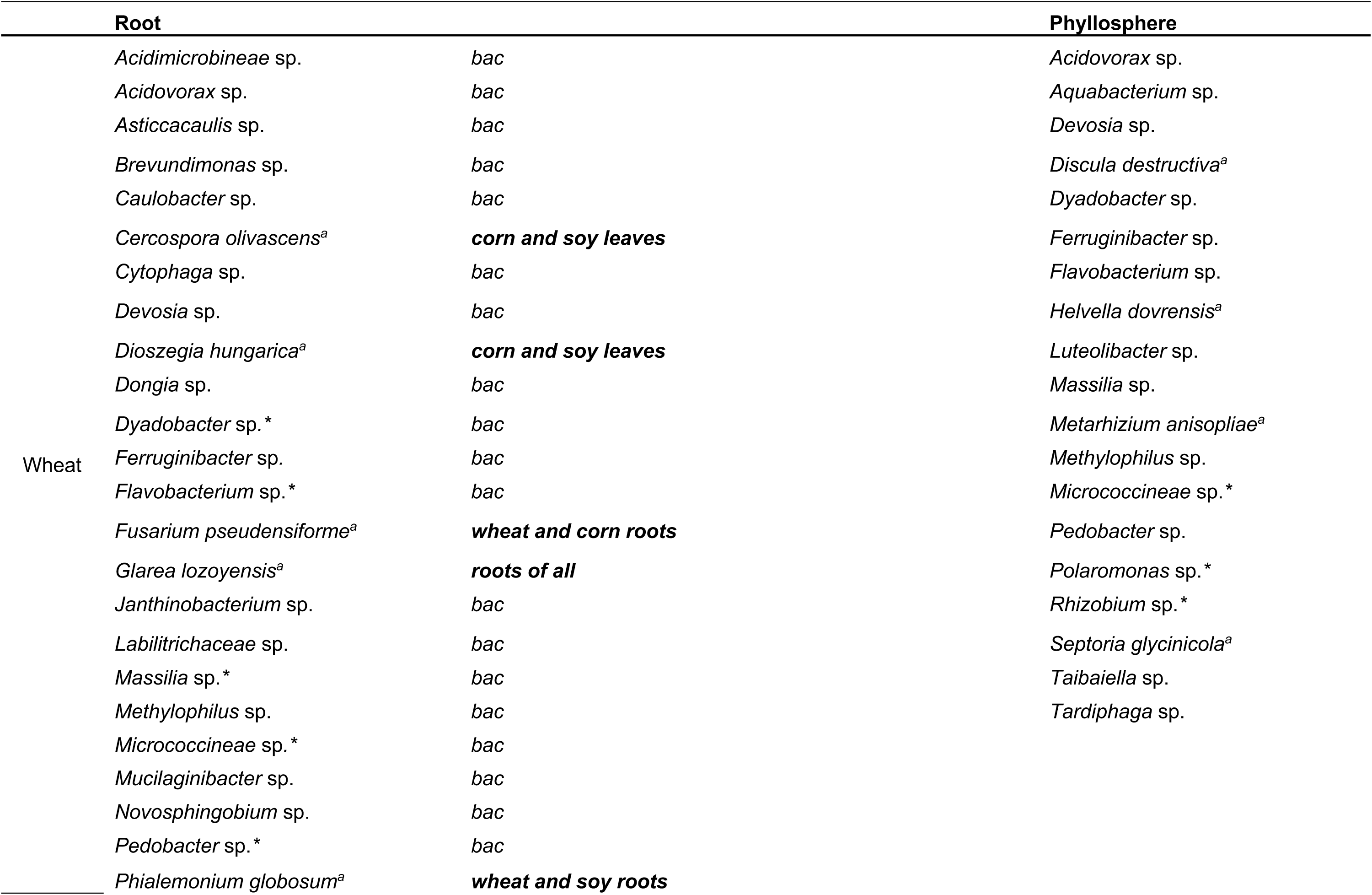
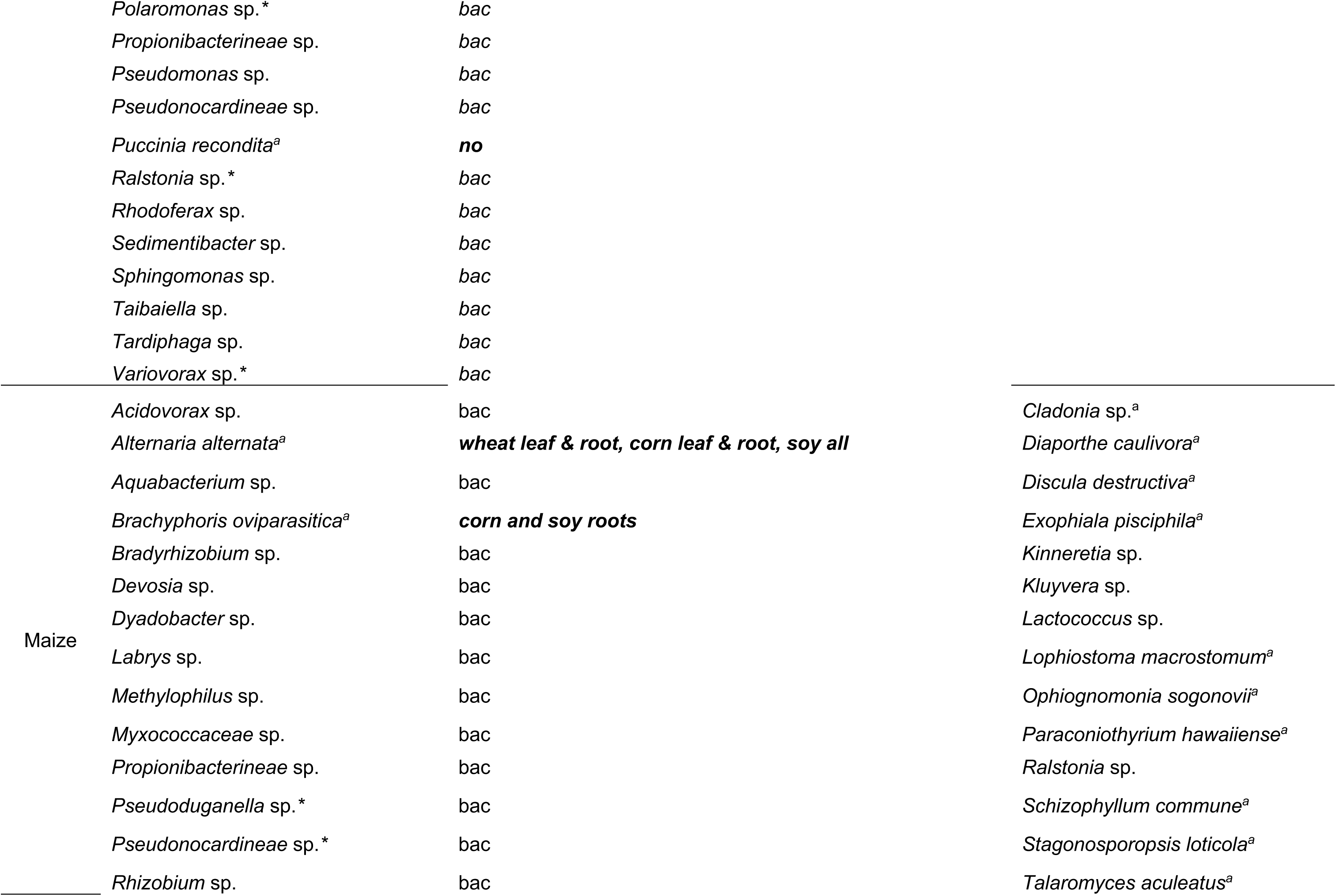
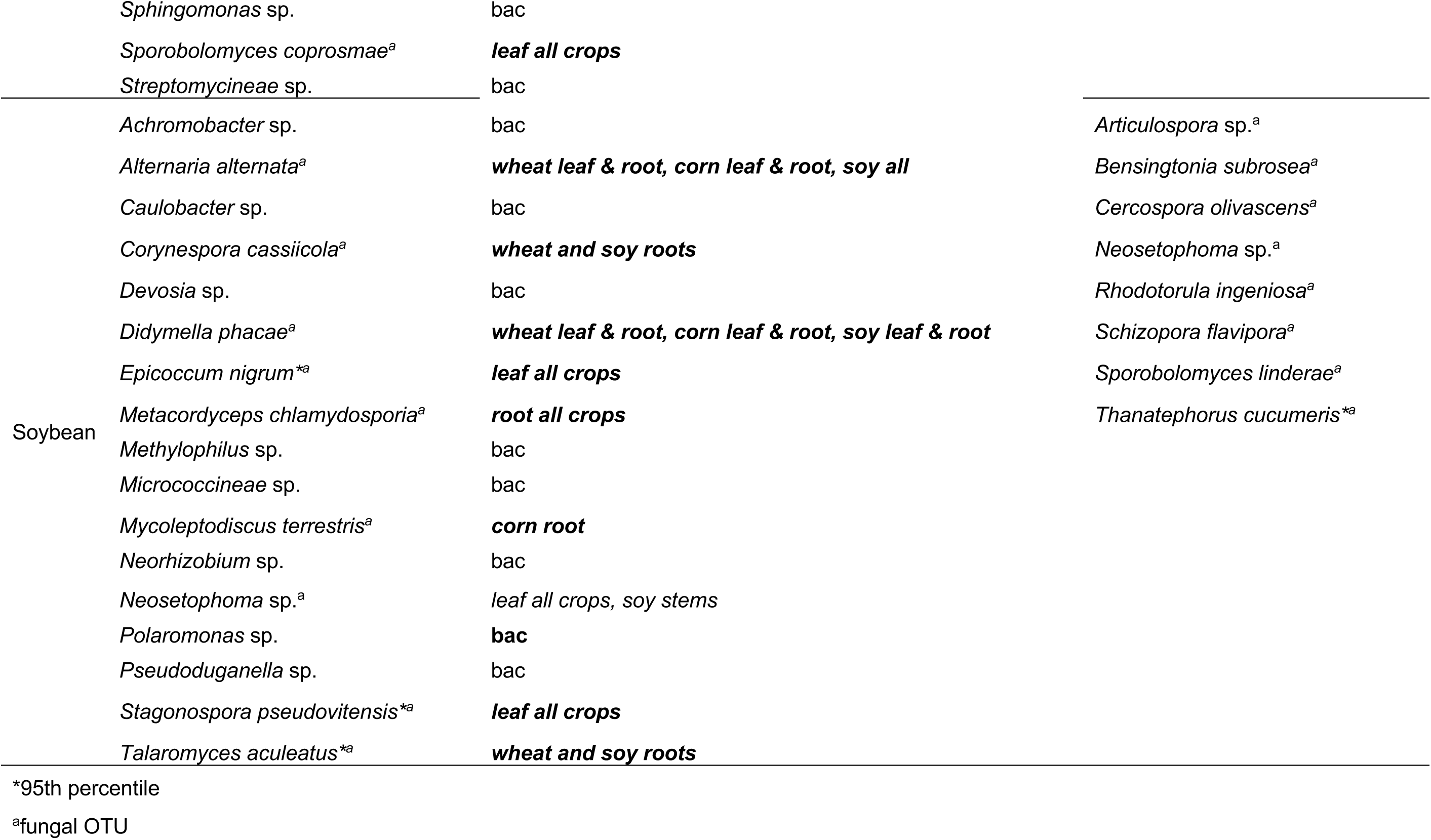
Hub taxa of KBS LTER crops.

**Figure 6.**
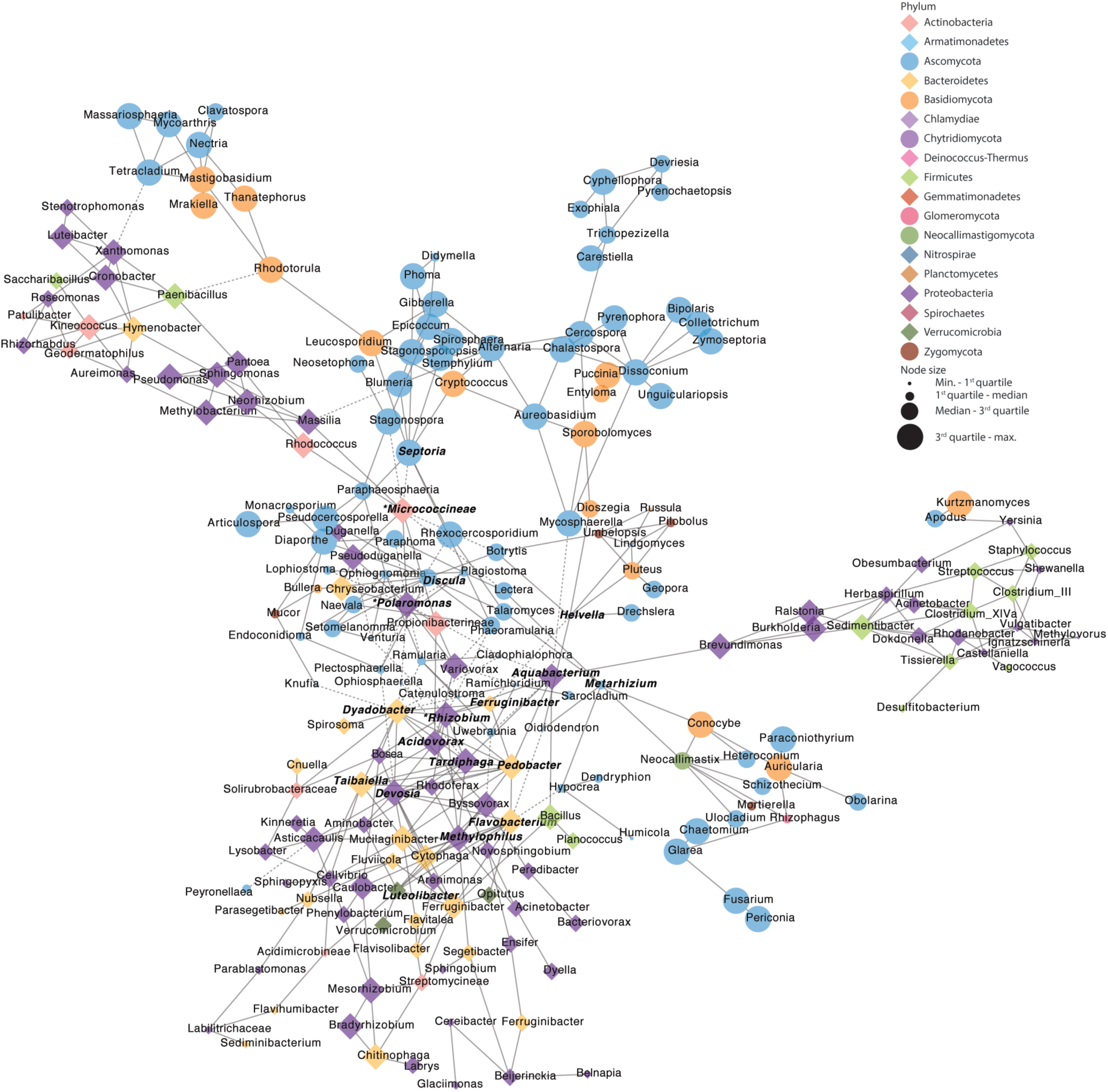
Co-occurrence network of taxa in wheat phyllosphere. Bacterial OTUs are represented by diamond shapes, fungal OTUs are represented by circles, and nodes are colored by class. Solid lines indicate positive correlation, and dashed lines indicate a negative correlation between OTUs. Node size indicates degree of connectivity. Node size indicates relative abundance, bolded node labels indicated hub taxa at the 75^th^ percentile, node labels marked with an asterisk indicate hub taxa at the 95^th^ percentile.

The bacterial hub taxa in maize roots were frequently positively correlated with other bacterial species. However, bacterial hubs had negative correlations with fungal nodes, with the exception of *Myxococcaceae* sp. and *Pseudonocardineae* sp., which were negatively correlated with each other (Fig. S12). The Glomeromycota, which form endomycorrhizal symbioses with plants, clustered in a small isolated network not connected with the main maize root network (Fig. S12). Hub taxa in the maize phyllosphere were largely identified as fungi, with only four hubs identified as bacteria (Fig. 7).

**Figure 7.**
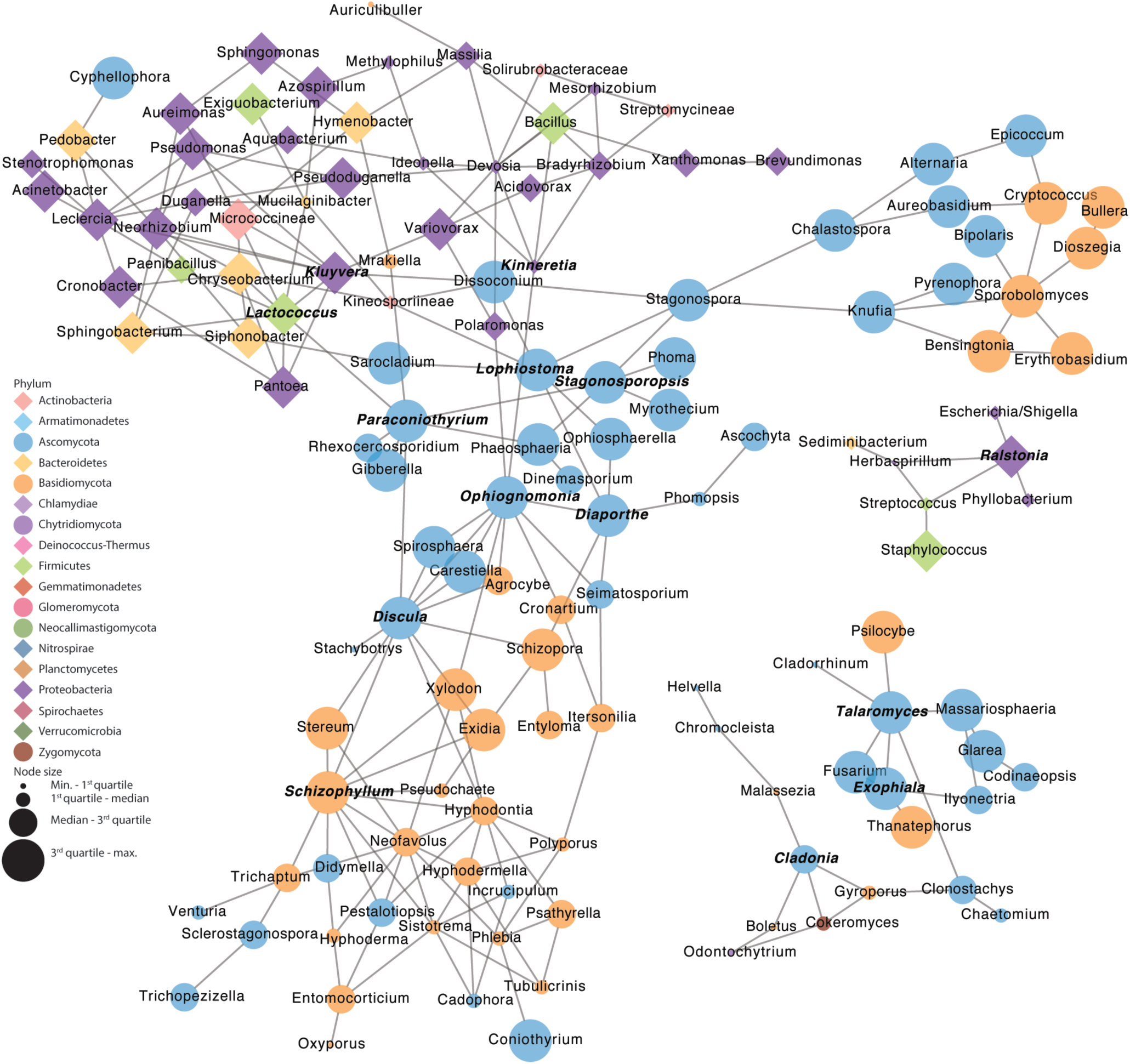
Co-occurrence network of taxa in maize phyllosphere. Bacterial OTUs are represented by diamond shapes, fungal OTUs are represented by circles, and nodes are colored by class. Solid lines indicate positive correlation, and dashed lines indicate a negative correlation between OTUs. Node size indicates degree of connectivity. Node size indicates relative abundance, bolded node labels indicated hub taxa at the 75^th^ percentile, node labels marked with an asterisk indicate hub taxa at the 95^th^ percentile.

Hub taxa were shared across plant organs, and they were also shared across host species. *Discula destructiva* was identified as a hub taxon in both wheat and maize phyllospheres, and was significantly associated with maize leaves as shown by indicator species analysis (Table S6). Four hub taxa were shared across maize and soybean roots (Table 2). Only one hub taxon, *Neosetophoma* sp., was shared between soybean root and phyllosphere networks (Figs. 8, S13) and was found to be significantly associated with soybean leaves (Table S6). Closely related taxa, such as several nodule-forming bacterial OTUs (*Bradyrhizobium, Mesorhizobium, Neorhizobium*) frequently occurred together in the networks. *Fusarium* sp. was present in all the co-occurrence networks and was identified as a hub taxon in the wheat phyllosphere.

**Figure 8.**
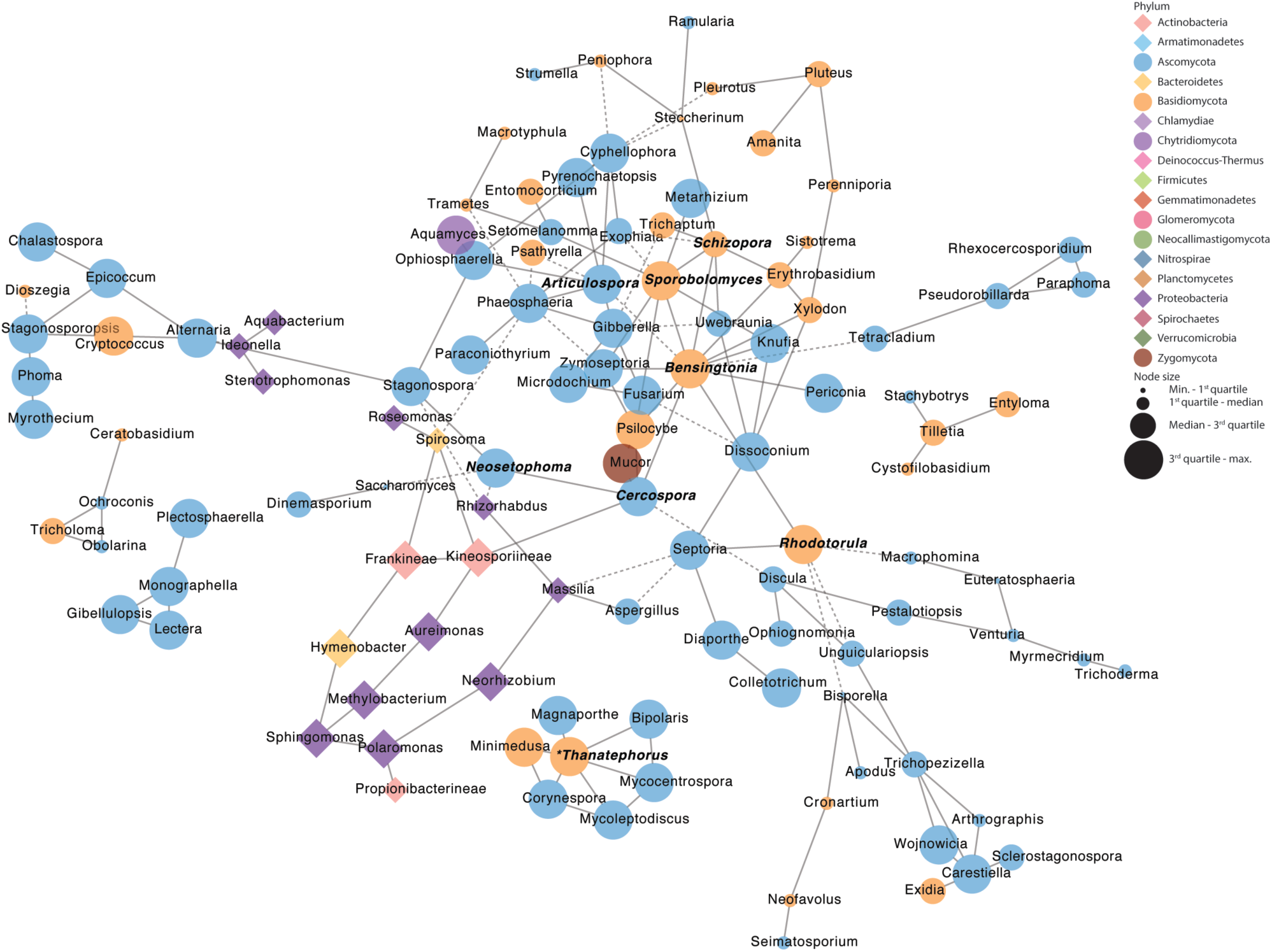
Co-occurrence network of taxa in soybean phyllosphere. Bacterial OTUs are represented by diamond shapes, fungal OTUs are represented by circles, and nodes are colored by class. Solid lines indicate positive correlation, and dashed lines indicate a negative correlation between OTUs. Node size indicates degree of connectivity. Node size indicates relative abundance, bolded node labels indicated hub taxa at the 75^th^ percentile, node labels marked with an asterisk indicate hub taxa at the 95^th^ percentile.

## DISCUSSION

Here we present an in-depth comparative analysis of fungal and bacterial microbial communities that colonize the major US row crops (wheat, maize, and soybeans) across a three-year rotation. Although these plant hosts share microbial taxa, the communities of the individual crops also harbor distinct OTUs, as determined by core taxa analysis and diversity metrics. Global analysis revealed distinct community clusters for above-ground versus below-ground organs. In a composite analysis of all crops and treatments combined, we also demonstrated that a subset of fungi were specifically and uniquely associated with a single plant organ: leaf, stem, or root. These organ-specific taxa represent a small subset of the fungal microbiome, yet they provide some clues as to the mechanisms of assembly of the microbiome. For many fungi, the spores are the agents of dispersal, having evolved to optimally distribute propagules to target areas where they can thrive (Calhim et al, 2018; Deacon 1980; Evans et al. 2016; Fischer et al. 2010; Pringle et al. 2015; Pringle et al. 2016). Thus, we further examined these taxa in terms of characteristics that affect fungal propagule distribution: spore size and shape, pigmentation, and means of dispersal. From that analysis we see that the majority of taxa exhibiting organ specificity are associated with spore characteristics that would enhance dispersal to the target organs.

Modes of dispersal can be generalized among the spore types of the major taxa found uniquely on specific plant organs. In our analysis, differentials are greatest among fungi unique to above-ground versus below-ground organs. Root-associated arbuscular mycorrhizal fungi of the Glomeromycetes, are associated with very large, generally spherical spores (Chaudhary et al. 2020), that commonly remain in the soil where produced (Egan et al. 2014; Warner et al. 1987; Watkinson 2016). However, recent work has shown that several taxa of AM fungi have spore traits that permit aerial some dispersal, including smaller spores (Chaudary et al. 2020). Fungi with the spore traits “up to 10 µm” had several different categories of dispersal, with the largest numbers of OTUs in the categories “aerial, soil”, “hypogeous”, “insect, other”, and the “up to 10 µm, lichenized/aerial, soil”. This spore size appears to very much depend on dispersal mode for targeting location. Whereas those categories with “aerial, soil” dispersal demonstrated a higher presence in above ground versus below-ground tissues, and those OTUs in the “hypogeous” and “insect, other” categories showed higher presence in the roots. The physics of aerial dispersal and spore traits do support this mechanism (Calhim et al. 2018; Deacon 2006; Pringle et al. 2015; Trail et al. 2005; Trail 2007). Proof that the taxa we have described as organ specific were placed at these organs directly due to means of dispersal and spore traits would necessitate collecting propagules directly as they land and demonstrating they colonize specific tissues. However, their absence from other organs across three years and three host species supports the conclusion that the fungi are relying on these mechanisms for their distribution.

Fungi that dominate the phyllosphere, including leaf pathogens and endophytes, typically produce spores (10 to 100 µm) that are airborne and often melanized, to protect from UV damage and desiccation (Eisenmann & Casadevall 2012; reviewed in Henson et al. 1999 and Jacobson 2000). In our study, these include the rusts and Dothidomycetes. The rusts are notable leaf pathogens with spores known for very long-range dispersal (Brown & Hovmøller 2012; Golan & Pringle 2017). Dothidiomycetes, the largest class of Ascomycota, includes *Alternaria spp*. (pigmented airborne leaf-infecting fungi (Pastor & Guarro, 2008)), and the frequently occurring Dothidiomycetes orders Pleosporales and Capnodiales, (Haridas et al. 2020), with a significant majority of OTUs associated with leaves. The Agaricales are distributed between leaves and roots. Their distribution to roots as well as leaves may reflect their abundant distribution under the leaf canopy, with basidiocarps forming on the moist soil surface through colonization of rotting plant tissue, and firing spores using the requisite available moisture (Webster et al. 1984). The lower apparent distribution of this group to wheat may reflect differences in temporal microbial patterns, and the fullness of the canopy. Winter wheat was planted at this site, requiring fall plantings and overwintering of young plants with the first wheat collection months occurring before collection of the first growth stages of maize and soybeans. Further study is warranted to determine if the reduced colonization of wheat is due to host genetics or environmental factors.

Bacterial diversity has most often been studied in the roots (Morris et al. 2017), and our results show that the greatest reservoirs of bacterial diversity are the roots of soybeans (71% OTUs are root-specific) and maize (53% OTUs being root-specific). However, a larger proportion of bacterial OTUs are shared among the three organs in wheat (only 36% are root-specific). Many previous studies have indicated that bacteria mainly colonize through the roots and of these, a few continue to colonize upward to the phyllosphere (reviewed in Compant et al. 2010). There are several recent publications suggesting the importance of bacterial diversity in the soil for maintaining plant productivity (Chen et al. 2020b; Lau & Lennon 2012), and bacterial diversity in the phyllosphere has been shown to be essential for ecosystem productivity (Laforest-Lapointe et al. 2017), in which the endophytic population is in part nurtured by the plant (Chen et al. 2020a). Further research is needed to determine whether bacteria have different roles in wheat versus soybeans and maize based on which organ harbors the most diversity.

Our findings did not provide strong evidence that management strategies alter microbial diversity more than effects of plant organ or host species. Our data suggest that root associated microbial communities are recruited by the plant hosts and selectively colonize the endophytic compartment. In this study and others, the influence of management strategies on root-associated or root endophytic compartments of row crops is minimal and community differences at these sites are more strongly influenced by the host plant (Naylor et al. 2017). Studies that describe an influence of management on plant-associated microbiota of row crops have been confined to the rhizosphere or bulk soil (for example, Peiffer et al. 2013; Strom et al. 2019; Wattenburger et al. 2019). Additionally, the influence of management on phyllosphere microbiota has been found to be minimal in several studies (for example, Gdanetz & Trail 2017; Longley et al. 2020; Sapkota et al. 2015). Although not directly tested in this study, the patterns of organ- and host-specificity observed support selective colonization of the root endophytic compartment.

We demonstrate that root-associated microbial communities are distinct from those of the phyllosphere in the three-crop rotation at the KBS-LTER site. Studies that analyze microbial communities across multiple organs, tissues, or compartments on the same plant often show that there are distinct communities across these niches (Coleman-Derr et al. 2015; Gdanetz & Trail 2017; Longely et al. 2020; Ottesen et al. 2013; Strom et al. 2019). Yet, until very recently, studies analyzing soil and rhizosphere microbial communities (for example, Copeland et al. 2015; de Souza et al. 2016; Gomes et al. 2018; Rojas et al. 2017; Strom et al. 2019) have been far more common than studies of phyllosphere communities. Although the rhizosphere and roots have been found to have a higher abundance and diversity of plant-associated bacteria (present study; Longely et al. 2020) and fungi (Ottesen et al. 2013) compared to the phyllosphere, microbes inhabiting aboveground plant organs and compartments are also important to plant health (Remus-Emsermann & Schlechter 2018; Ryffel et al. 2015; Vorholt 2012). Research on model systems shows that phyllosphere microbes can influence disease occurrence and nutrient cycling (Abanda-Nkpwatt et al. 2006; Delmotte et al. 2009; Vorholt 2012) and some microbes have a life cycle that is not captured by a soil- or root-associated phase, for example, important plant pathogens such as rust fungi (Webster & Weber 2007).

Although in this study the conventional, no-till, and low input managements included fungicide- and insecticide-treated seed, there were no controls to assess the direct impact of fungicide treatments on the seed microbiome. Noel et al. (2020) did not detect a difference in oomycete composition in the soybean seedling rhizosphere with or without anti-oomycete chemical seed treatments. Nettles et al. (2016) have demonstrated that fungicide seed treatments affect the rhizosphere fungal community of maize and soybean but had differential effects on the phyllosphere as tested during the vegetative growth phase. Therefore, the effect of fungicide seed treatments on the microbiome may be contextual. Studies in mature plants have not been reported. Surprisingly little information is available on the effects of seed treatments on non-pathogenic fungi in the microbiome (Lamichhane et al. 2020). This is an area worthy of further exploration.

Our findings indicate that fungal diversity in maize and soybeans decrease across the growing season, while bacterial diversity remains constant in these two plant hosts. Some aspects of microbial diversity, specifically the alpha-diversity described in this study, appear to vary temporally across the seasons, but this may also be an artifact of seasonal changes such as temperature and moisture (Lauber et al. 2013). Previous studies examining changes in plant-associated microbial communities in a variety of crops over the course of a growing season have not identified a consistent trend. Some studies found increasing community diversity over time (in apples (Shade et al. 2013) and winter wheat (Gdanetz & Trail 2017)) whereas decreases in diversity have been documented in soybeans and sugarcane (Copeland et al. 2015; Rascovan et al. 2016), and no significant changes in spring wheat (Sapkota et al. 2017). Additional studies are needed to examine how temporal plant microbial community dynamics of row crops are being influenced by maturation (and likely senescence) of the host plant and environmental factors, such as intensity and duration of light, temperature, and moisture. Environmental changes are likely to affect individual microbial species differently. Computer modelling of the plant microbiome suggests that initial colonizers influence final community structure (Evans et al. 2016). Further investigations should be conducted to experimentally determine the effect of random colonization by microbes on the composition of the community at important plant life stages. An understanding of the mechanisms by which the initial colonizers influence final community composition will have implications for community manipulation in both field crops and synthetic environments such as greenhouses or hydroponic systems.

Hub taxa are important members of the plant-associated microbiota that can influence final community composition and plant health (Agler et al. 2016). In the current study, putative hub taxa identified in the plant organs of one host species were not always shared across all others. However, these taxa are commonly occurring OTUs found across all hosts and management strategies. In addition, the fungal hub species included pathogens of other plant species. For example, *Discula destructiva*, a hub species of both the wheat and maize phyllosphere, causes dogwood anthracnose. In previous studies, the presence of non-host pathogen hubs have been shown to prevent the infection by another pathogen (Agler et al. 2016; Chen et al. 2018) perhaps via induced resistance. As observed in this study and previously described (Berry & Widder 2014), closely related species designated by OTUs of Rhizobia, wood rot fungi, and mycorrhizal fungi frequently co-occur in network analyses. Also demonstrated here and shown with inferred networks (Agler et al. 2016), microorganisms interact with each other across Kingdoms and thus it is important to examine cross-kingdom interactions. Hub taxa identified in this study, with few exceptions, were also identified as significantly associated with a specific plant organ by indicator analysis, coupled with their specific means of distribution which target them to that organ. These organisms warrant further study to determine their roles in the microbiomes of healthy plants. Confirmation of hub taxa and understanding of their influence on community composition will be important for community-scale microbial manipulation to improve agriculture, including in precision agricultural approaches (Tackenberg et al. 2016).

This study shows that plant organs (leaves, stems, and roots) have distinct microbial communities. Plant-associated microbial communities arise from the microbial reservoir of the surrounding ecosystem, but for fungi, spore traits maximize their ability to target specific organs. Assembly of microbial communities has been shown to also be affected by host genetics (Chen et al. 2020a; Hassani et al. 2018; Maignien et al. 2014). Research has indicated that the host can selectively manipulate colonization by some microbes and this selection may be in the form of seed or root exudates that recruit microbes for better (Wu et al. 2015), or for worse (Nelson 2004). Host genotype has also been shown to have a role in phyllosphere mycobiome (Sapkota 2015), and in particular, leaf chemistry has been shown to suppress spore germination (Barilli et al. 2016). While host genetics can be at least partially controlled by breeding, intentional manipulation of the microbiome is less straightforward. A recent study shows that in addition to host genotype, the order of species arrival affects the phyllosphere fungal community outcome (Leopold & Busby 2020). As we show here, fungi evolve to distribute where they will thrive, so dispersal, also plays a role in developing the fungal community. This suggests that those species with adaptations that allow them early targeted arrival will have more success in predominating in their host plants. Only a single year was examined for each host, and additional years are in progress to examine the formation of the microbiomes in the three-crop rotation to better resolve factors involved in organ specific communities and the effect of individual hosts. Our long-term goal is to the formation and stability of the microbial communities under the influence of climate change.

## Acknowledgments

We thank Mitchell Roth for assistance in collecting soybean samples. The authors acknowledge the support of the MSU Plant Sciences Fellowship Program, Michigan State University AgBioResearch, and the NSF Long-term Ecological Research Program (DEB 1637653) at the Kellogg Biological Station. This work was supported in part by NIFA grant MICL08541 from the USDA National Institute of Food and Agriculture to FT.

## Author Contributions

FT and KG designed the experiments. KG managed the field collections, sample preparation and performed the data analysis. ZAN conducted ternary analysis and interpretation and participated in the writing and editing of the manuscript. KG and FT interpreted the data and wrote the manuscript.

## Figure Legends & Table Titles

**Table S1**. Gene primer sequences used in this study.

**Table S2**. Collection dates for plant growth stages.

**Table S3**. Sequence processing and OTU assignment statistics.

**Table S4**. Samples with zero high-quality reads

**Table S5**. Significantly different OTUs (Bonferroni adjusted p-value) identified with DeSeq2 across management strategies or growth stages of each host species.

**Table S6**. Propagule characteristics of fungal OTUs significantly associated with a host and/or organ.

**Table S7**. Mean estimated Shannon Diversity Index (*H’*) and standard deviation.

**Table S8**. Mean observed OTUs and standard deviation. Superscripts indicate significant differences within a growth stage and management style, after analysis of variance and Tukey’s honest significant difference test.

**Table S9**. Permutational multivariate analysis of variance (PERMANOVA) results of Bray-Curtis values. Analysis is illustrated in Figure 5.

---

**Figure S1.**
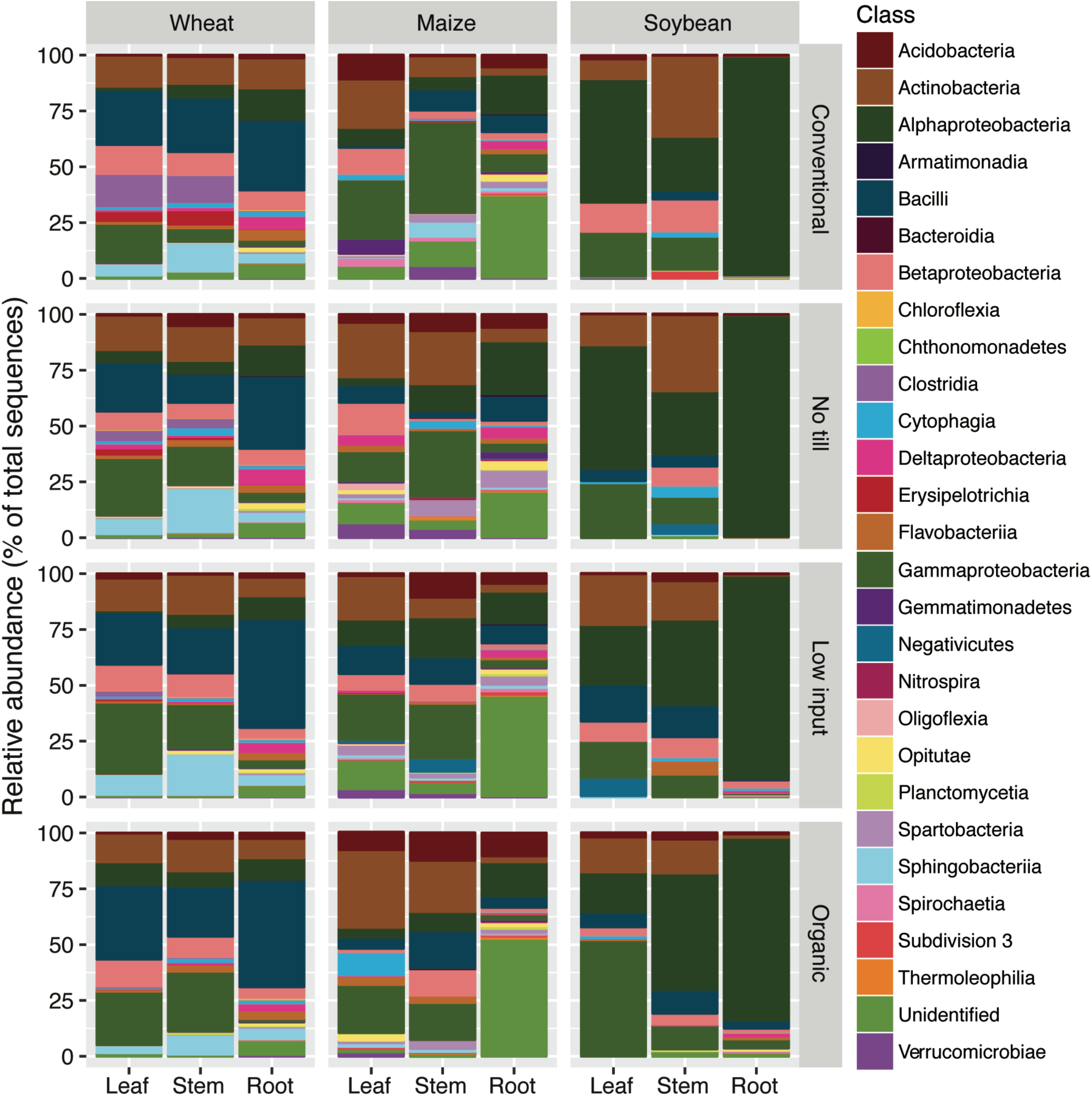
Class-level relative abundance of OTUs in bacterial communities across crop, organ, and management strategies.

**Figure S2.**
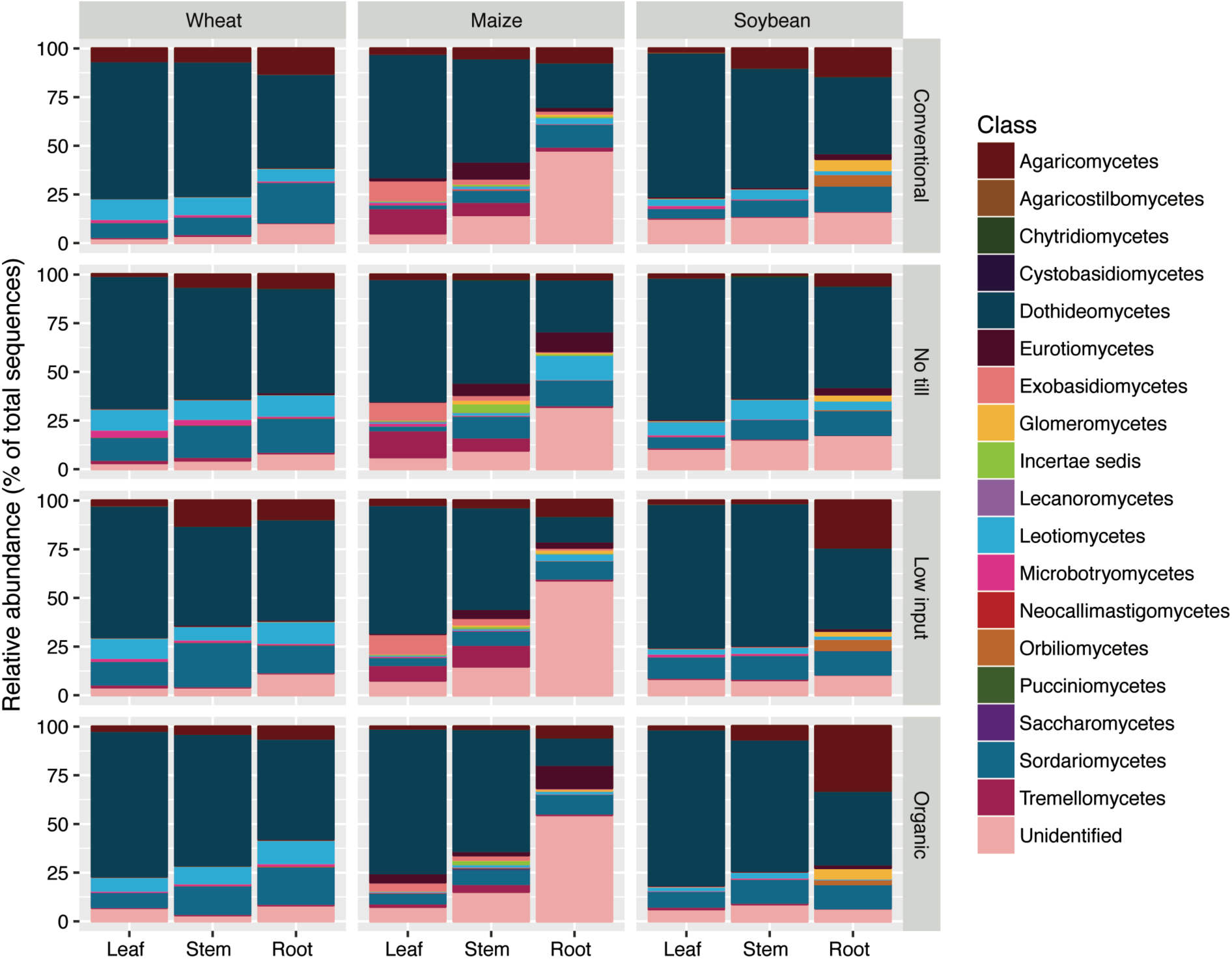
Class-level relative abundance of OTUs in fungal communities across host, organ, and management strategies.

**Figure S3.**
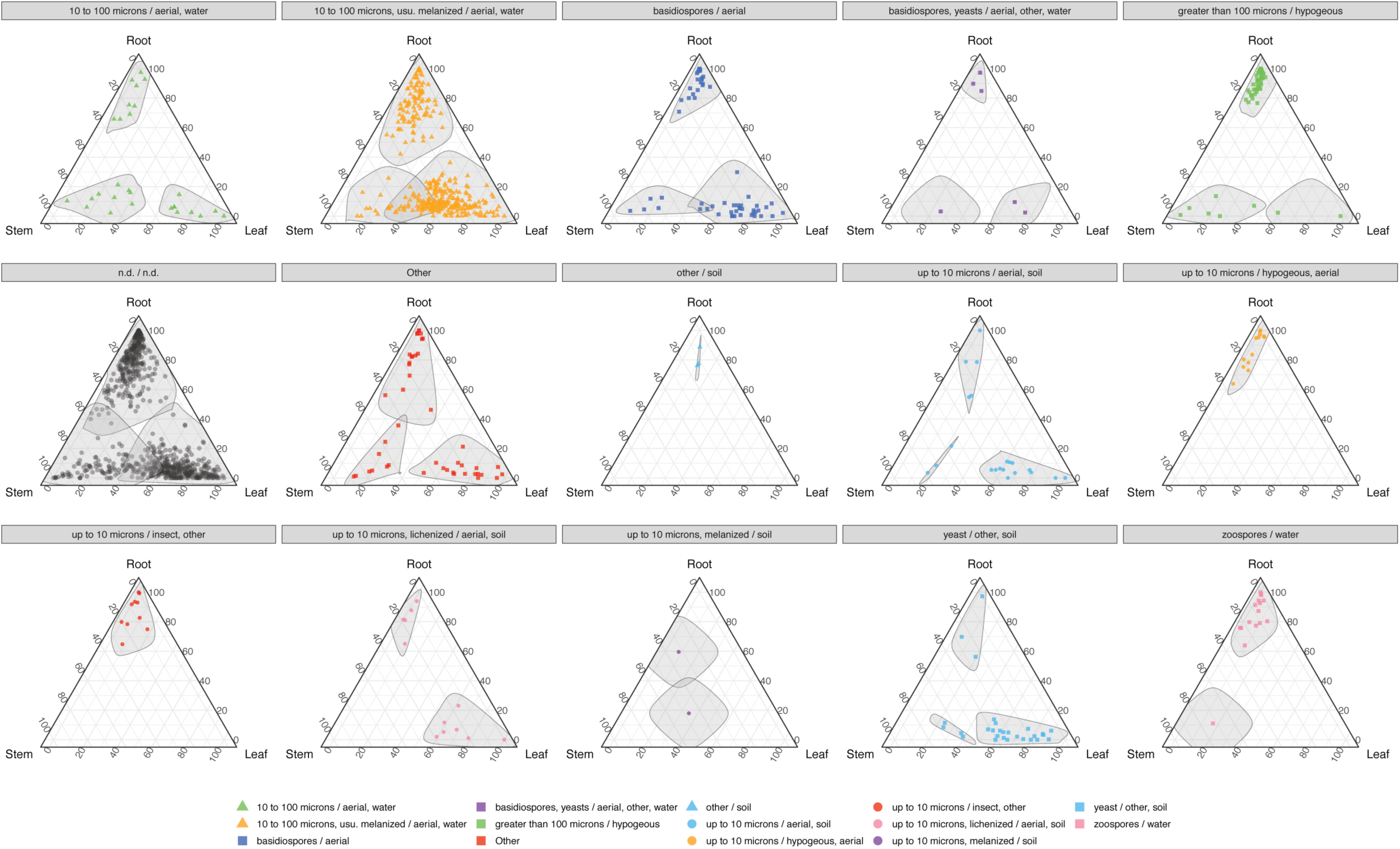
The distribution of propagule characteristics across plant organs. Colored shapes representing propagule characteristics were significant indicators of plant organs with a Benjamini-Hochberg corrected P-value (α = 0.05). Shaded areas encircle OTUs that are significant indicators for a plant organ. Grey points were significantly associated with an organ, but the propagule characteristics could not be determined (n.d. / n.d.).

**Figure S4.**
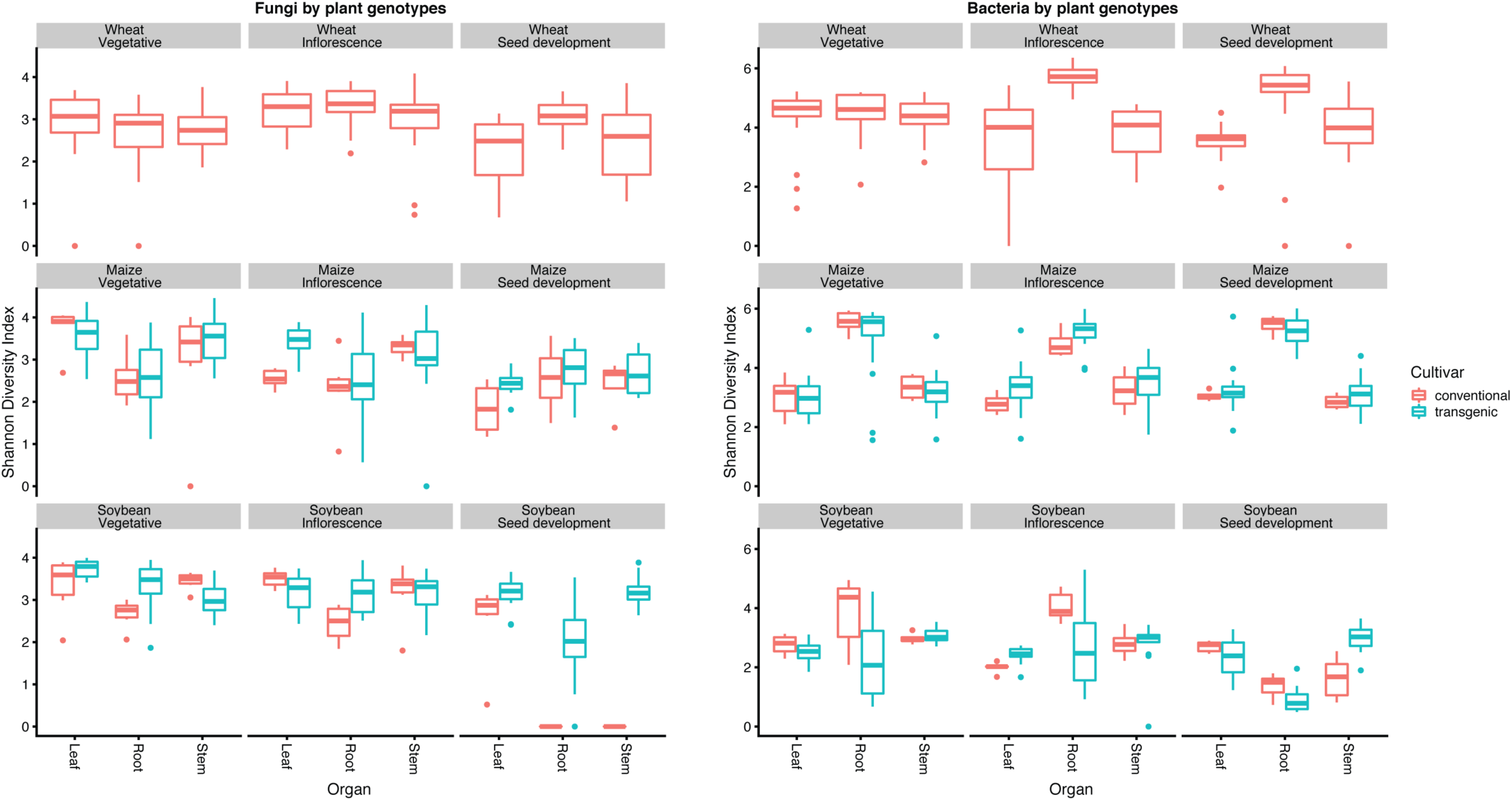
Alpha diversity of management strategies with (T1 = conventional, T2 = no-till, T3 = low inputs in maize and soybeans) and without (T1-T4 wheat, T4 = organic in maize and soybeans) transgenic cultivars; fungi (left panels) and bacteria (right panels). Comparisons are made between Shannon Diversity Index (H’) of conventional or transgenic cultivars (maize and soybeans only).

**Figure S5.**
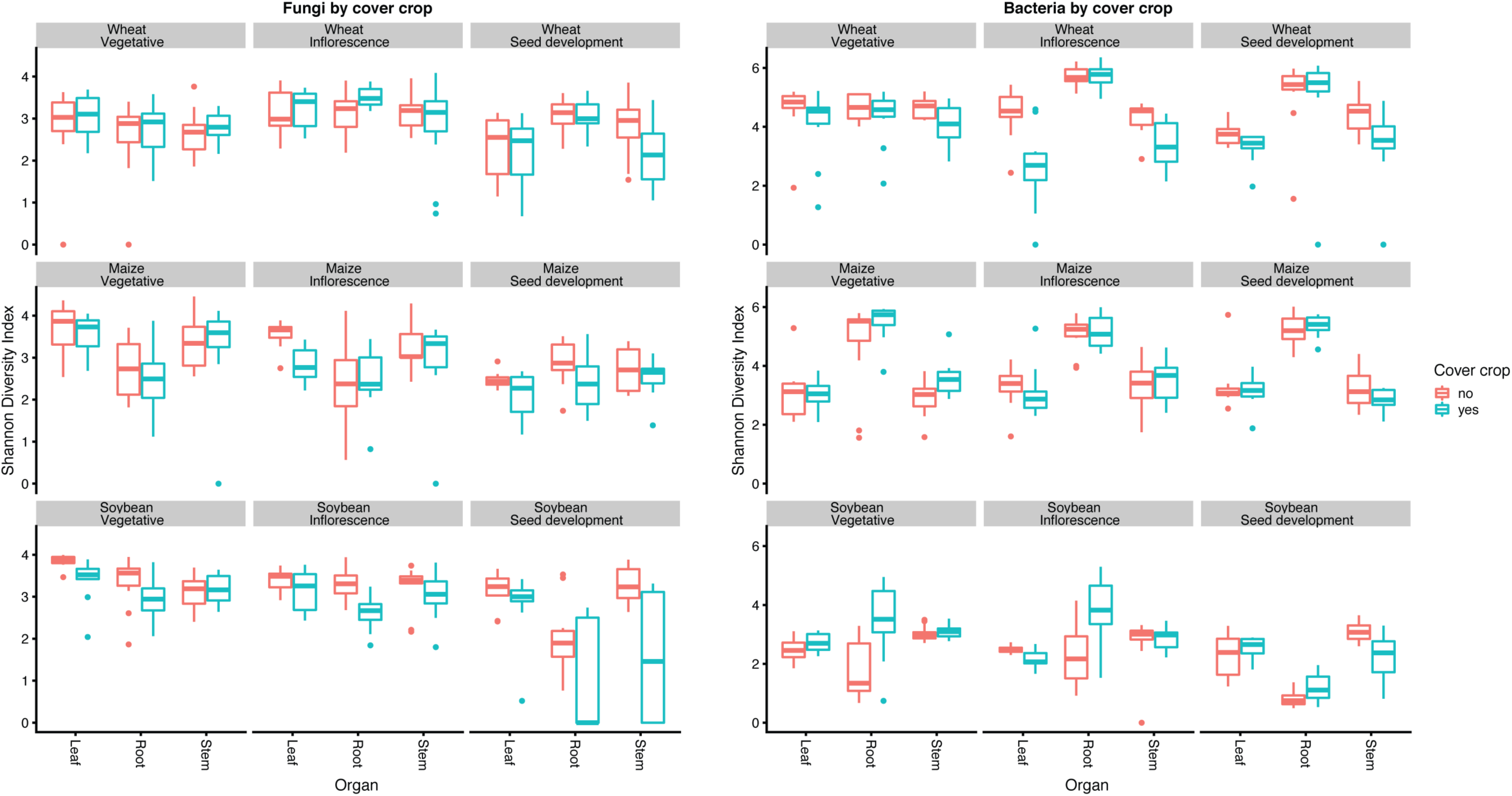
Alpha diversity of management strategies with (low inputs and organic) and without (conventional and no-till) cover crops; fungi (left panels) and bacteria (right panels). Comparisons were made within Shannon Diversity Index (H’). Microbial communities inhabiting each of the organs (leaves, stems, or roots) were compared for plots with and without cover crops within each collection period and host plant species.

**Figure S6.**
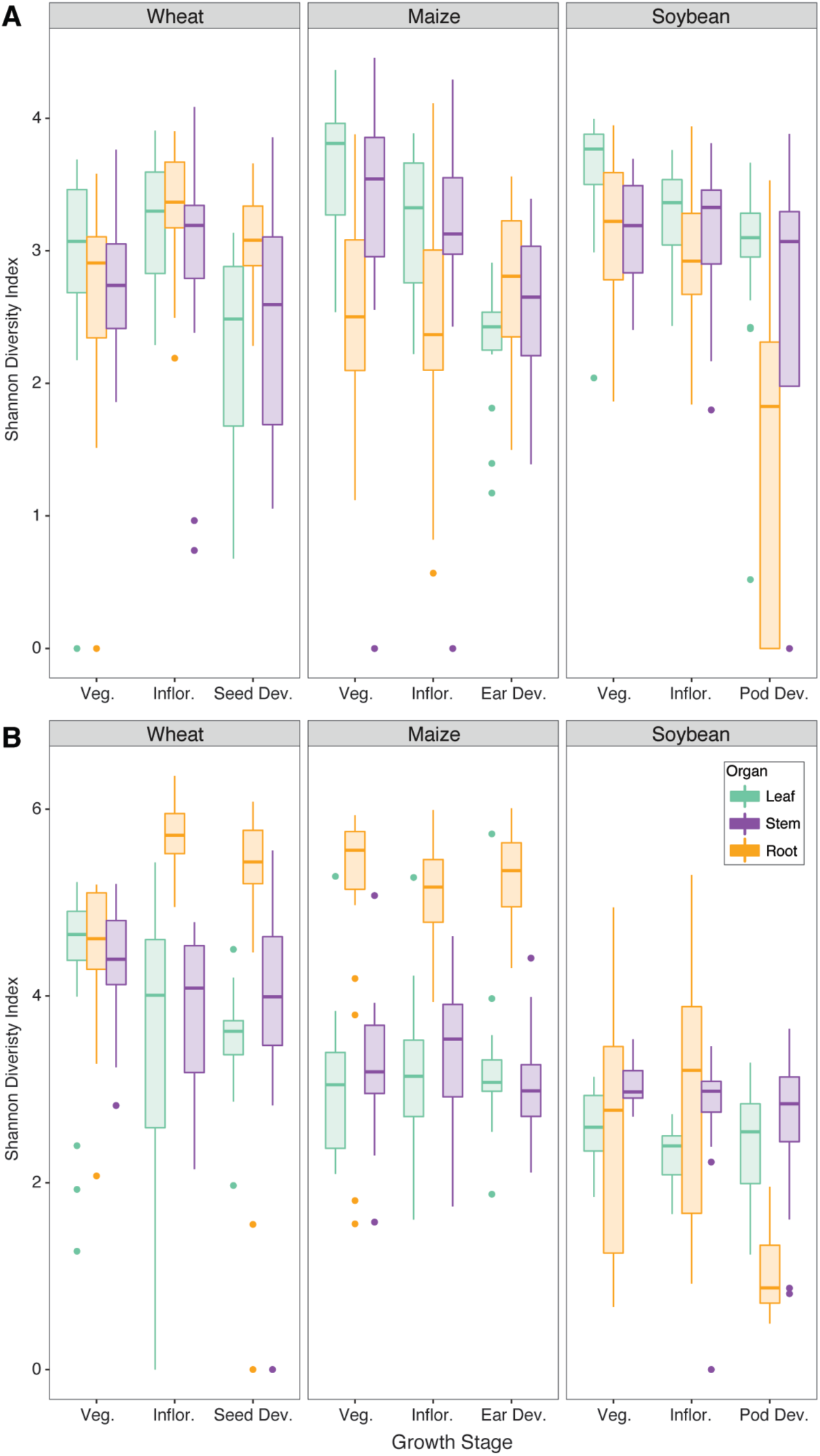
Global alpha diversity patterns calculated by Shannon Diversity Index (H’). All treatments and replicates were pooled within growth stages for (A) fungi and (B) bacteria. Data are represented by 24 replicates from each host species-growth stage-organ combination. Center line of boxes represents the median of samples. The upper and lower sides of the boxes represent the third and first quartiles, respectively. Whiskers represent ± 1.5 times the interquartile range. Data points beyond whiskers represent outliers. Analysis of variance and Tukey’s honest significant difference were used to test significance (P < 0.05). Statistical support is detailed in Table S7.

**Figure S7.**
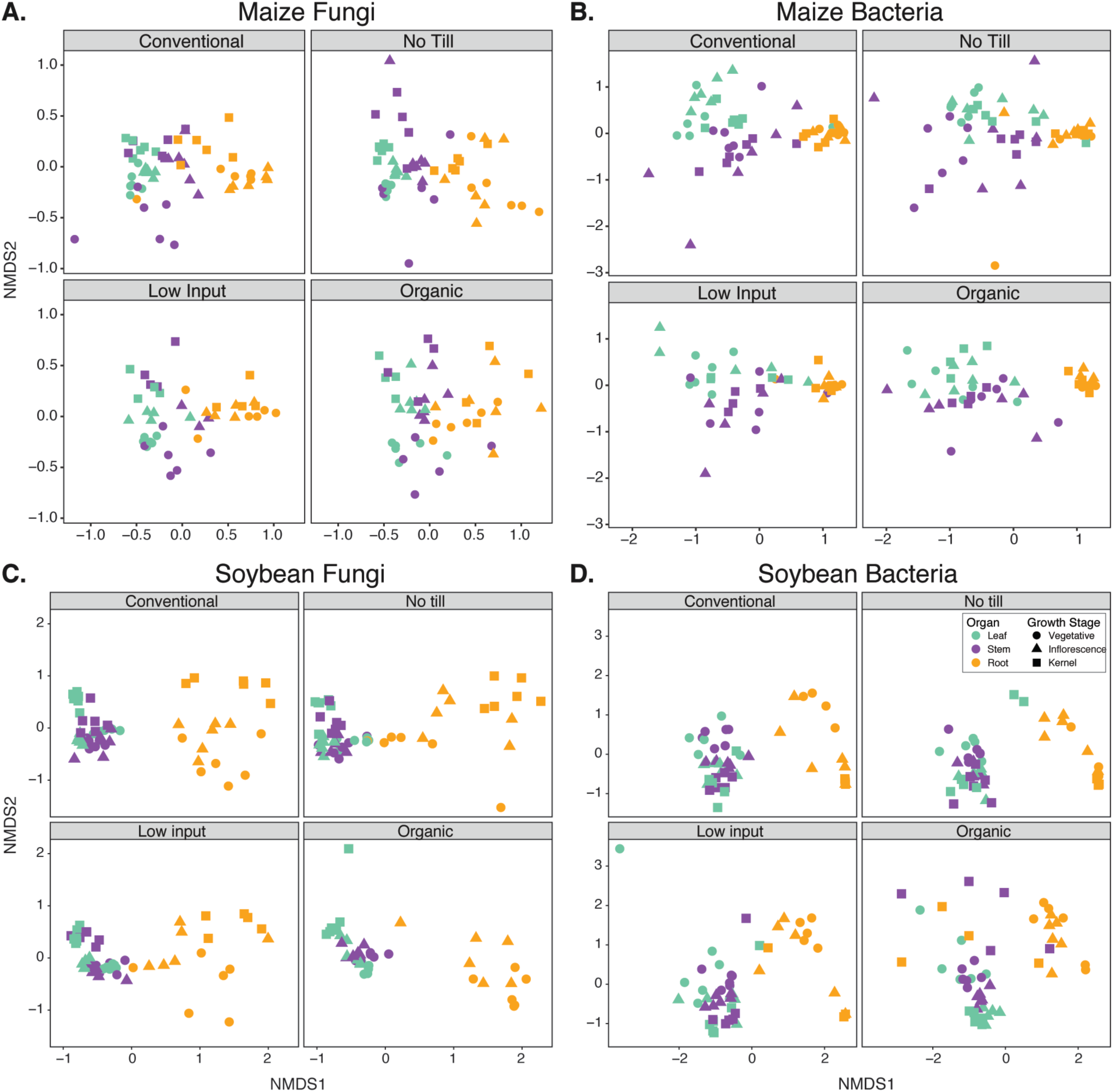
Influence of management strategies on beta diversity of (A) maize fungal, (B) maize bacterial, (C) soybean fungal, (D) soybean bacterial communities originating from each plant organ. Non-metric multidimensional scaling (NMDS) calculated by Bray-Curtis distances. Statistical support is detailed in Table S9.

**Figure S8.**
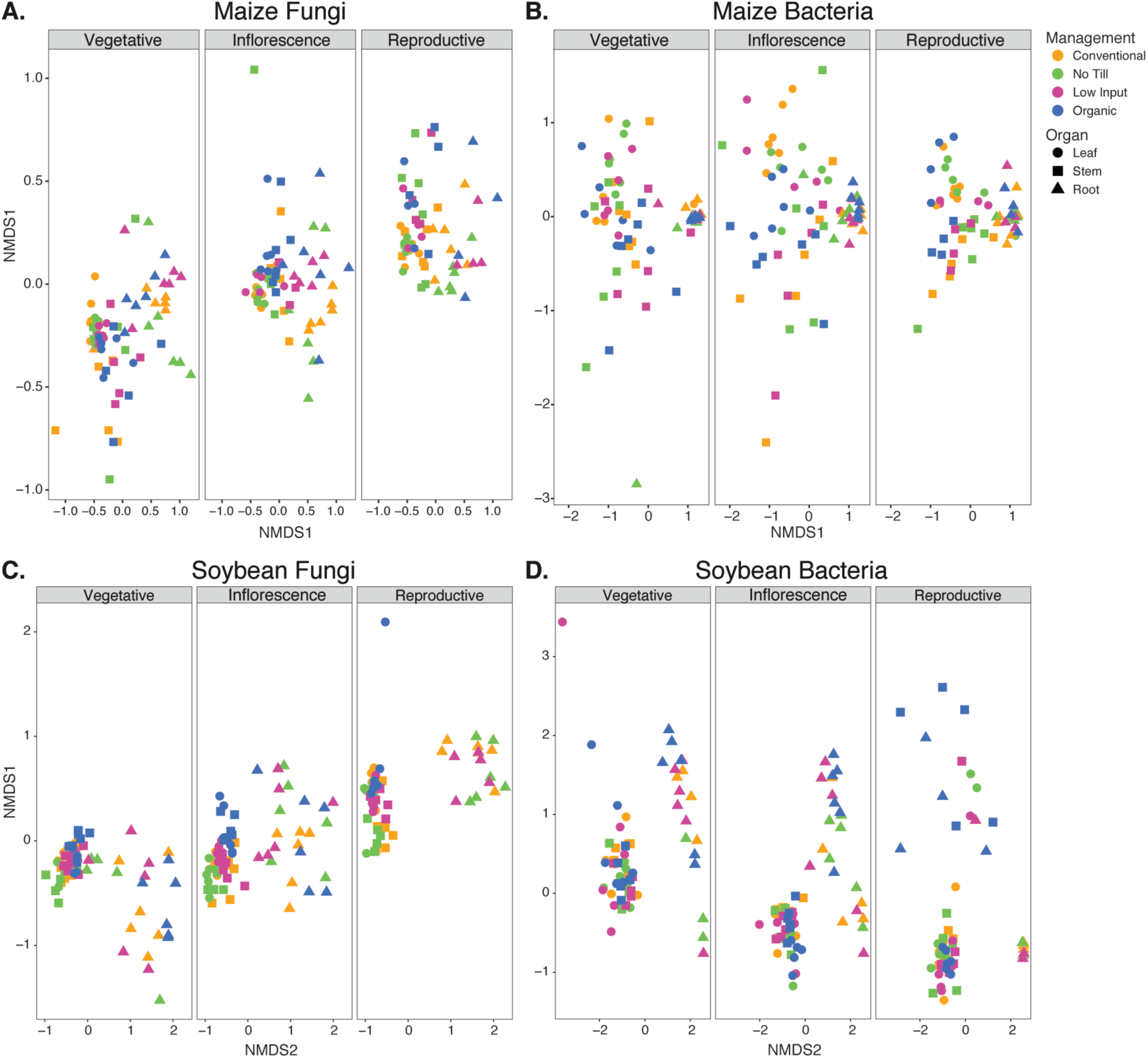
Effect of growth stage on beta diversity of (A) maize fungal, (B) maize bacterial, (C) soybean fungal, (D) soybean bacterial communities originating from each management strategy. Non-metric multidimensional scaling (NMDS) calculated by Bray-Curtis distances. Statistical support is detailed in Table S9.

**Figure S9.**
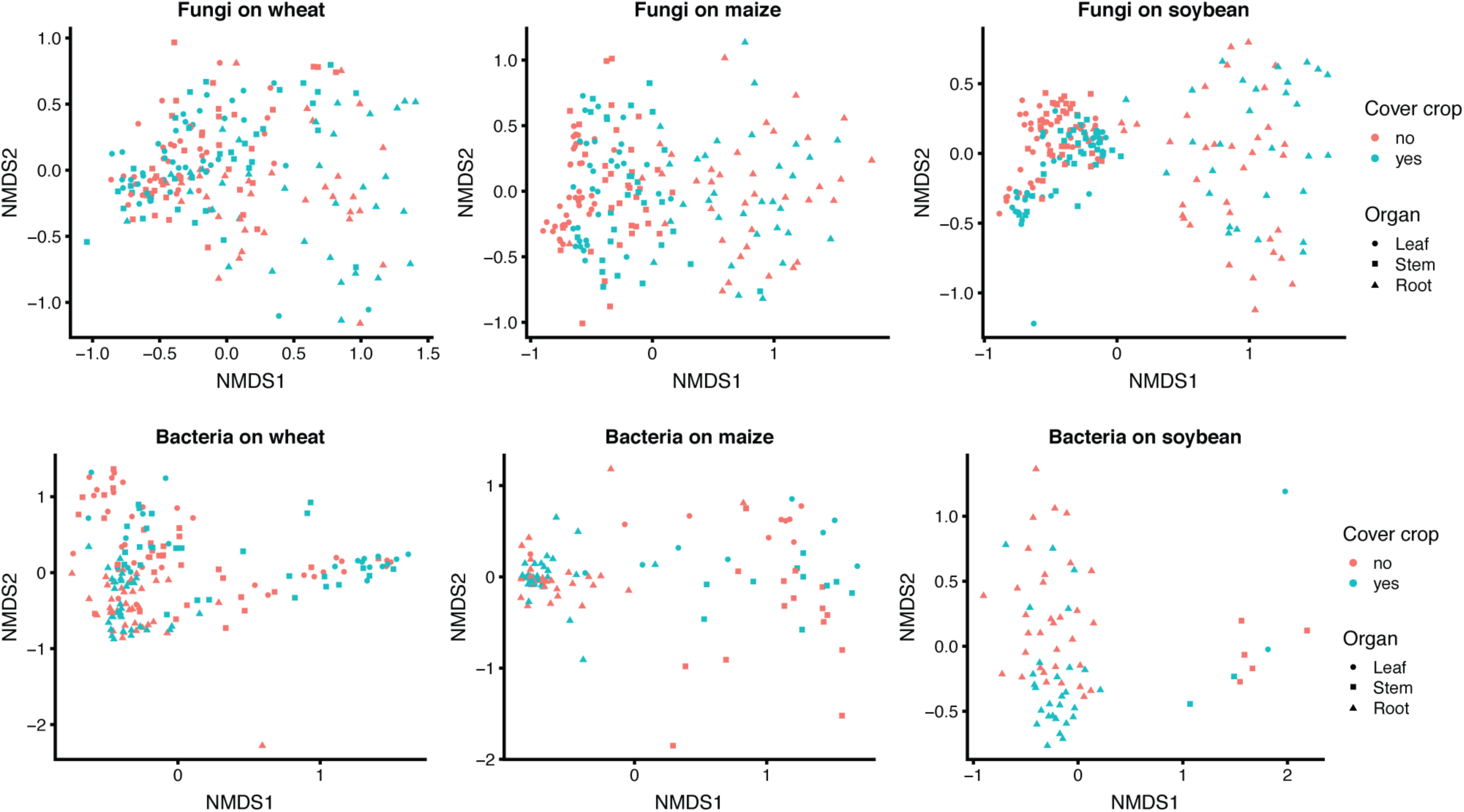
Effect of cover crop on beta diversity of fungal (upper row) and bacterial (lower row) communities. Low input (T3) and organic (T4) managements planted with red clover cover crops, and conventional (T1) and no-till (T2) received no cover crop. NMDS calculated using Bray-Curtis distances. Statistical support is detailed in Table S9.

**Figure S10.**
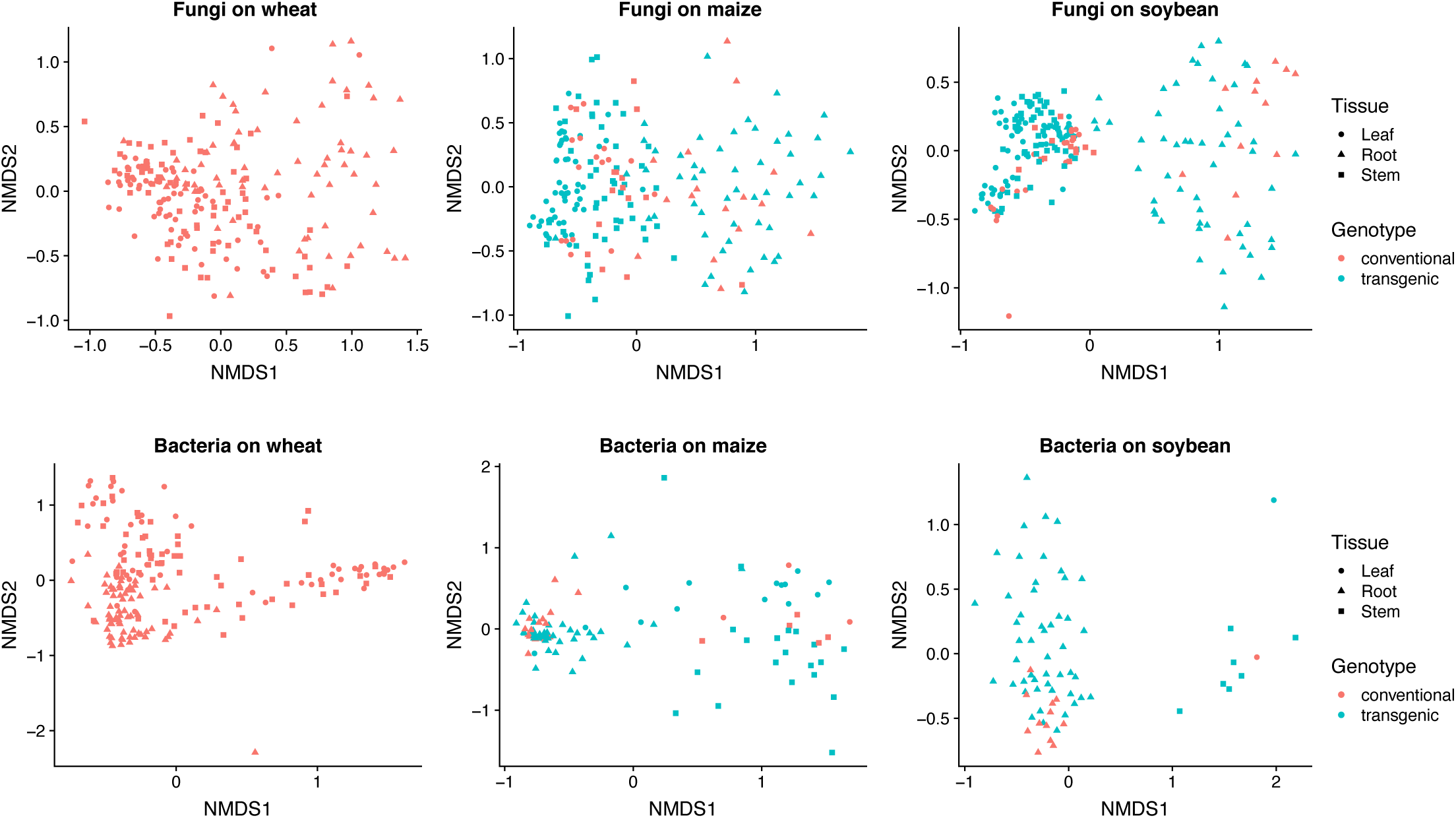
Effect of cultivar on beta diversity of fungal (upper row) and bacterial (lower row) communities. Non-metric multidimensional scaling (NMDS) calculated by Bray-Curtis distance. Maize and soybeans in T1 = conventional, T2 = no-till, T3 = low inputs were transgenic cultivars, and all wheat (T1-T4), T4 = organic maize, T4 and soybeans were planted with conventional cultivars. Statistical support is detailed in Table S9.

**Figure S11.**
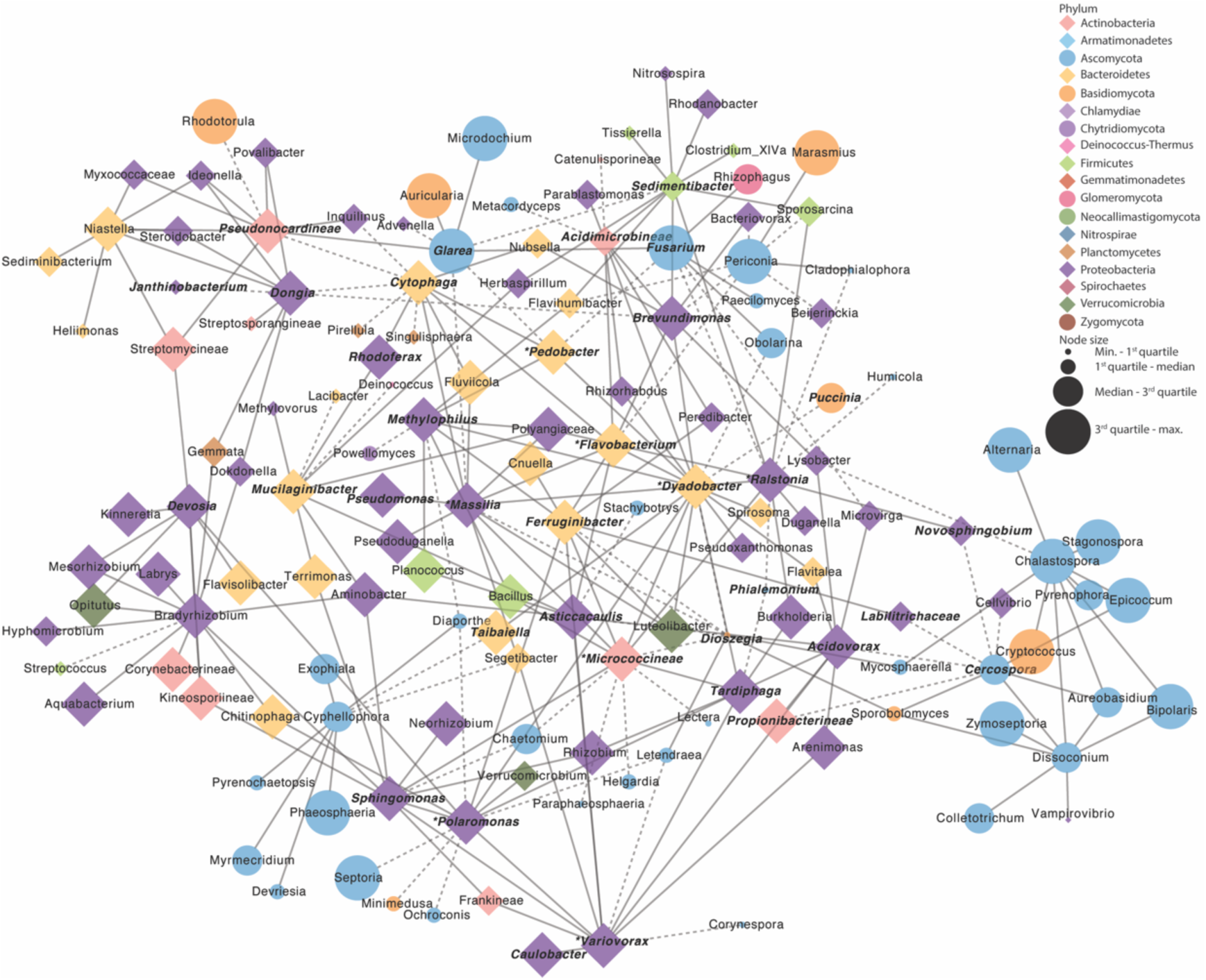
Co-occurrence network of taxa in wheat roots. Bacterial OTUs are represented by diamond shapes, fungal OTUs are represented by circles, and nodes are colored by class. Solid lines indicate a positive correlation, and dashed lines indicate a negative correlation between OTUs. Node size indicates relative abundance, bolded node labels indicate hub taxa at the 75th percentile, node labels marked with an asterisk indicate hub taxa at the 95th percentile.

**Figure S12.**
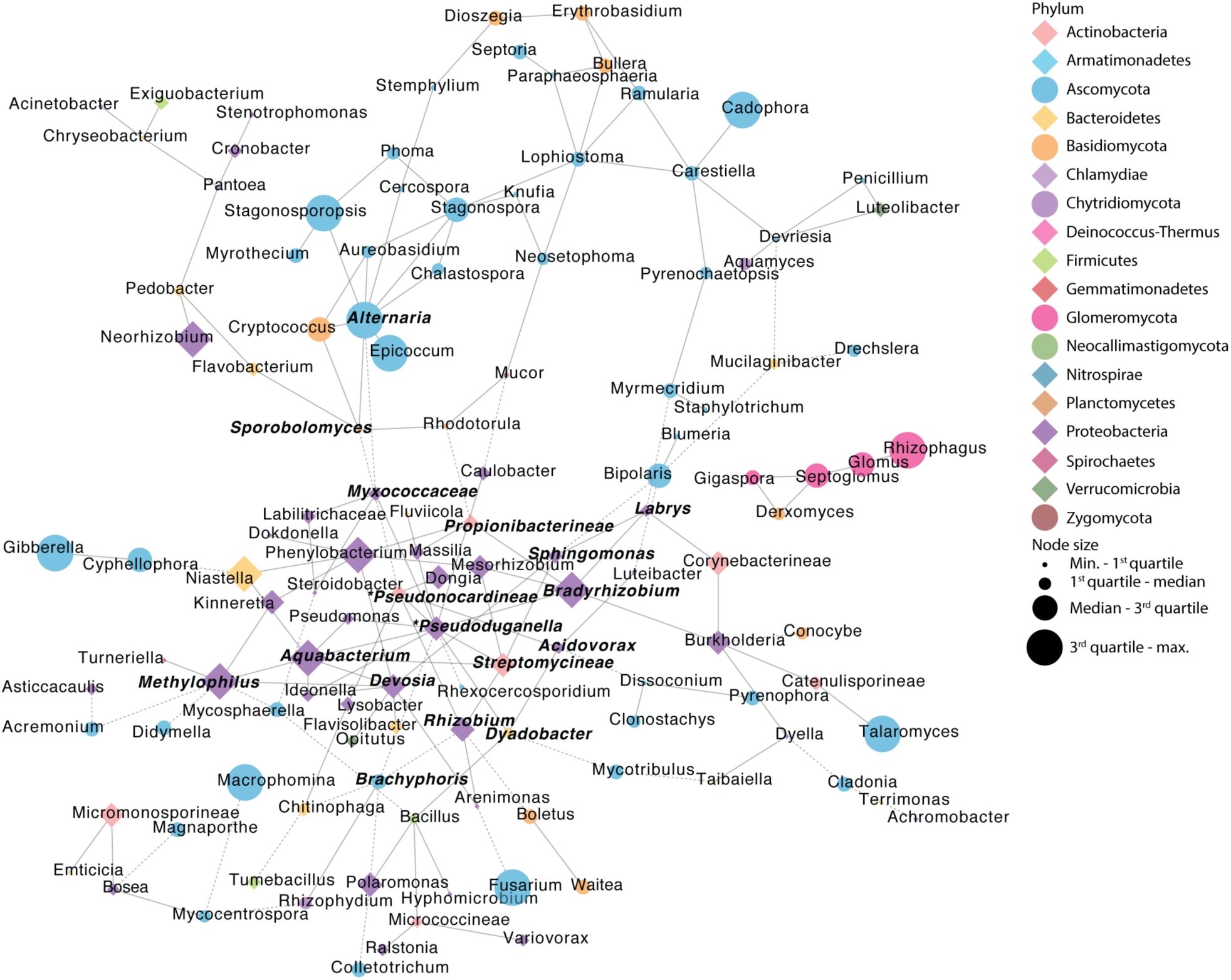
Co-occurrence network of taxa in maize roots. Bacterial OTUs are represented by diamond shapes, fungal OTUs are represented by circles, and nodes are colored by class. Solid lines indicate positive correlation, and dashed lines indicate a negative correlation between OTUs. Node size indicates relative abundance, bolded node labels indicate hub taxa at the 75th percentile, node labels marked with an asterisk indicate hub taxa at the 95th percentile.

**Figure S13.**
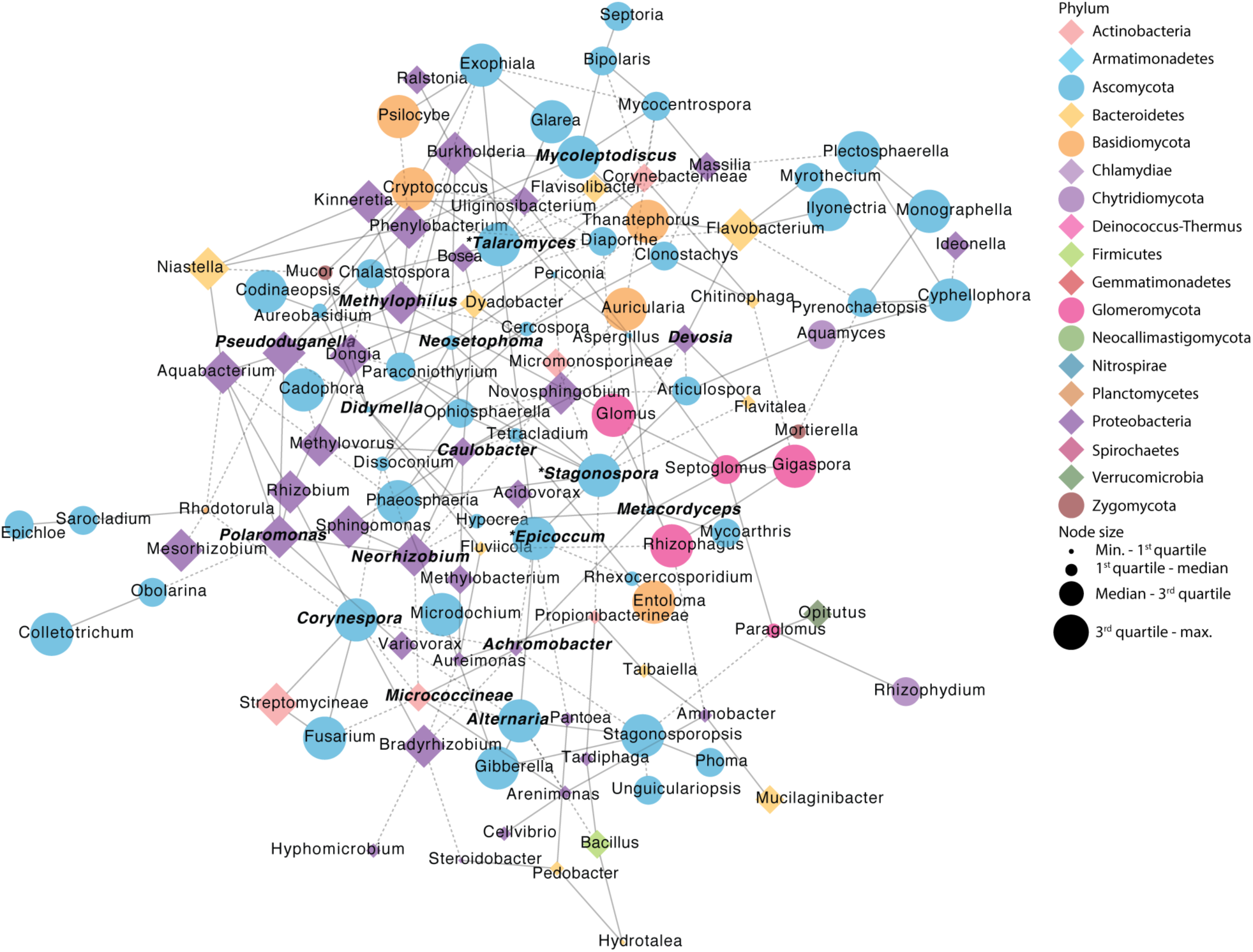
Co-occurrence network of taxa in soybean roots. Bacterial OTUs are represented by diamond shapes, fungal OTUs are represented by circles, and nodes are colored by class. Solid lines indicate positive correlation, and dashed lines indicate a negative correlation between OTUs. Node size indicates relative abundance, bolded node labels indicate hub taxa at the 75th percentile, node labels marked with an asterisk indicate hub taxa at the 95th percentile.

